# GM-CSF instructs vertebral emergency myelopoiesis in neuropathic pain

**DOI:** 10.1101/2025.11.23.690022

**Authors:** Glauce R. Pigatto, Gabriel V. Lucena-Silva, William A. Gonçalves, Andreza U. Quadros, Samara Damasceno, Conceição E. A. Silva, Volodya Hovhannisyan, Tusar K. Acharya, Daniele C. Nascimento, Sara E. Jager, Carlos G. Meschick, Maria C. Cavallini, João P. Mesquita, Gabriel N. R. Goldstein, Adam J. Dourson, Helder I. Nakaya, James R. F. Hockley, Christian D. Ellson, Marcello H. Nogueira-Barbosa, Gileade P. Freitas, Marcio M. Beloti, Maíza M. Antunes, Gustavo B. Menezes, Ishwarya Sankaranarayanan, Khadijah Mazhar, Edward C. Emery, Kenneth L. Madsen, Jose C. Alves-Filho, Fernando Q. Cunha, Ari B. Molofsky, Anna V. Molofsky, Theodore J. Price, Julia E. Smith, Peter M. Grace, Burkhard Becher, Arkady Khoutorsky, Thiago M. Cunha

## Abstract

Meningeal immune cells have recently emerged as critical modulators of neural function in both physiological and pathological states^1,2^. In particular, immune cells within the meninges surrounding the dorsal root ganglia (DRG) have been implicated in the pathogenesis of neuropathic pain^3–5^. Yet, the cellular complexity of the meningeal immune landscape and the neuroimmune interactions linking this niche to neuropathic pain remain poorly understood. Here, we show that peripheral nerve injury induces both the recruitment of leukocytes to the DRG meninges and their transcriptional reprogramming across innate (primarily myeloid cells) and adaptive immune compartments. The accumulation of myeloid cells, predominantly neutrophils and classical monocytes, within the DRG meninges originates from local vertebral bone marrow (BM) emergency myelopoiesis, followed by their migration through ossified vertebral channels. Further analysis identified meningeal granulocyte-macrophage colony- stimulating factor (GM-CSF), produced mainly by group 2 innate lymphoid cells (ILC2s), as a key mediator that instructs vertebral BM emergency myelopoiesis after peripheral nerve injury, thereby promoting neuropathic pain. These findings uncover a fundamental process linking meningeal immunity to vertebral BM emergency myelopoiesis in the pathophysiological cascade of neuropathic pain and highlight meningeal GM-CSF as the instructive signal orchestrating this neuroimmune axis.

## Main

Chronic neuropathic pain is a consequence of injury or disease affecting the somatosensory nervous system, and its initiation and maintenance are strongly shaped by neuro-immune interactions spanning the peripheral and central nervous systems^6,7,8,9,10^.

Meningeal immune cells have been implicated in central nervous system (CNS) development, homeostasis, and various pathologies^1,11,12^. These cells produce bioactive molecules, particularly cytokines, that affect CNS function^13,14,15^. Conversely, meningeal immune populations can also sense the physiological and pathological states of the underlying parenchyma^16,17^. For instance, autoimmunity, injury, or neurodegeneration within the CNS can trigger immune responses that affect all meningeal layers^18,19,20,21^.

The meninges not only envelop the brain and spinal cord but, within the spinal column, the dura and arachnoid mater also extend laterally toward the DRG, forming a pocket-like structure around the proximal region of each ganglion^22^

The subdural space continues along the ganglia, where the dural layers of the CNS transition into the epi- and perineural structures associated with the DRG^22^. This intimate anatomical association might implicate meningeal immune cells in the surveillance of the DRG microenvironment and, in turn, these cells may secrete factors that modulate the function of DRG parenchymal cells and drive pain hypersensitivity.

Here, we sought to define the immune landscape of the DRG meninges, mapping its cellular composition, origins, and signaling circuitry, and to determine how these meningeal immune cells shape the development of neuropathic pain. We found that following peripheral nerve injury, GM-CSF, produced primarily by meningeal ILC2s, stimulates emergency myelopoiesis in the vertebral BM. This process drives the migration of myeloid cells through ossified vertebral channels toward the DRG meninges, where they promote pain hypersensitivity. Notably, blockade of GM-CSF alone was sufficient to disrupt this cascade and abolish neuropathic pain, revealing a previously unrecognized meningeal immunity–to– vertebral BM axis underlying pain development.

## DRG meningeal immunity in neuropathic pain

To assess changes in the immune landscape of the DRG meninges during peripheral nerve injury-induced neuropathic pain, we used the spared nerve injury (SNI) model^23^ (Fig. 1a). SNI induced pain hypersensitivity (Fig. 1b) and concurrently increased the number of leukocytes in the ipsilateral lumbar (L3-L5) DRG meninges (Fig. 1c; Extended Data Fig. 1a), peaking 10-14 days after injury (Fig. 1c). Leukocyte number in DRG meningeal tissue remained increased for up to 28 days and returned to basal levels 60 days after SNI (Extended Data Fig. 1b,c). This response was localized to the ipsilateral lumbar DRGs, as leukocyte numbers remained unchanged in DRG meninges from both the uninjured contralateral side (Fig. 1d) and the brain (Fig. 1e). Notably, leukocyte accumulation in DRG meninges after SNI did not differ between male and female mice (Fig. 1f). Immune cells also accumulated in the DRG meninges after chronic constriction injury (CCI), another model of neuropathic pain^24^ (Extended Data Fig. 1d).

**Fig. 1.**
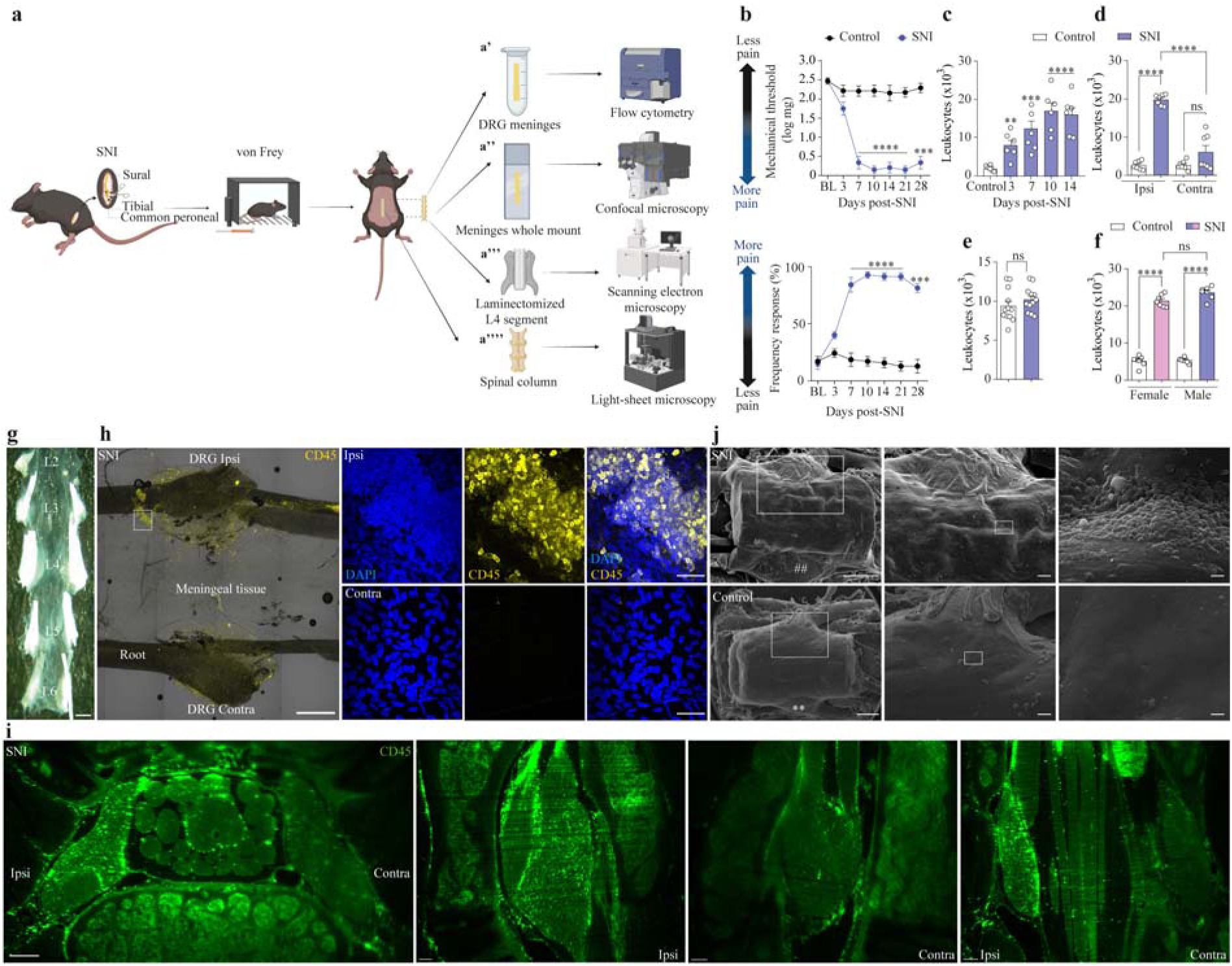
Peripheral nerve injury promotes leukocyte accumulation in DRG meninges. **a.** Schematic illustration of induction of spared nerve injury (SNI) model of neuropathic pain in C57Bl/6 mice followed by pain behaviors’ analyses (von Frey test) and surgery to isolate: **a’.** DRG meninges for flow cytometry; **a’’.** whole spinal meninges with DRGs attached for confocal analyses; **a’’’.** whole preparation of laminectomized spinal/DRG L4 segment for SEM; **a’’’’.** cleared spinal column for light-sheet microscopy. **b.** Time course of mechanical pain hypersensitivity evaluated before and at indicated times (days) after SNI. *n* = 8 per group. **c.** Absolute number of leukocytes (CD45^+^ cells) in the DRG meninges in control mice or at indicated times (days) after SNI. *n* = 6 per group. **d.** Absolute number of leukocytes (CD45^+^ cells) in the DRG meninges of control mice or 10 days after SNI ipsilateral (Ipsi) *versus* contralateral (Contra) sides related to the surgery. *n* = 8 per group. **e.** Absolute number leukocytes (CD45^+^ cells) in the brain meninges of control mice or 10 days after SNI. *n* = 12 per group. **f.** Absolute number of leukocytes (CD45^+^ cells) in the DRG meninges from male *versus f*emale mice control or 10 days after SNI. *n* = 7 per group. **g.** Overview of mouse lumbar meninges whole mount sample (from L2-L6 DRG levels) imaged by stereomicroscopy. Scale bar: 100 μm. **h.** Left panel: Representative maximum intensity projections of the mouse lumbar meningeal whole mount sample collected 14 days after SNI surgery. Low magnification view of L4 meningeal sample segment. Scale bar, 500 μm. Right panels: High magnification images were obtained at L4 DRG meninges level in the contralateral (Contra, bottom) or ipsilateral side (Ipsi, top). Samples were stained for CD45^+^ cells (yellow) and DAPI (blue). Scale bars, 50 μm. **i.** Representative images from high- resolution cleared-based light-sheet microscopy of spinal region of mice after SNI. Ipsi indicates the ipsilateral L4 DRG and CL indicates the contralateral side, respectively. Samples were stained for CD45^+^ cells (green) Scale bars: 100 μm (left panel) and 200 μm (middle and right panels). **j.** Scanning electron microscopy showing the dura mater outer surface shaped by subdural cells. Samples were collected from control or after SNI induction. Inserts show the ipsilateral region (left panels) selected for successive magnifications (middle and right). # indicates the ipsilateral L4 DRG and ## the contralateral L4 DRG, respectively. Scale bars (from left to right), 500, 100, and 10 μm. Statistical analysis: two-way ANOVA with Bonferroni post-test (b); one-way ANOVA with Tukey post-tests (c,d,f), unpaired two-sided *t*-tests (e). \*\**P* < 0.01, \*\*\**P* < 0.001, \*\*\*\**P* < 0.0001; ns= non-significant. *n* = biologically independent samples from individual mice or mouse tissues. Error bars indicate the mean ± s.e.m. Illustrations were created with BioRender.com (https://biorender.com).

Next, using a whole-mount tissue preparation containing the lumbar spinal meninges together with the DRG and their associated meninges (Fig. 1g), we confirmed a robust increase in leukocytes (using immunostaining for CD45^+^ cells) ipsilateral to the nerve injury (Fig. 1h; Extended Data Fig. 1e,f). Leukocytes accumulated primarily in the meninges surrounding the L4 and L5 DRGs, which contain most of the sensory neuron cell bodies forming the injured nerve, but were also found in the L3 and L6 DRG meninges (Extended Data Fig. 1g). To extend these observations, light-sheet 3D microscopy of the cleared spinal column revealed a pronounced increase in leukocytes post-SNI in the dorsal and ventral regions of the ipsilateral L4 and L5 DRG meninges (Fig. 1i, movies 1-3). The increased presence of cells beneath the dura mater after SNI was further evident in scanning electron microscopy (SEM) images of the spinal segment containing the L4 DRG (Fig. 1j). Together, these findings indicate that peripheral nerve injury induces localized leukocyte accumulation in the ipsilateral DRG meninges.

To assess the specific subpopulations of DRG meningeal leukocytes and their responses to nerve injury, we performed single-cell RNA sequencing (scRNA-seq) of leukocytes isolated from the DRG meninges (Fig. 2a). Clustering analysis of transcriptomes from 8,538 DRG meningeal leukocytes (SNI: 4,925 cells; Control: 3,613 cells) revealed substantial differences between SNI and control mice (Fig. 2b). Overall, we identified 11 main clusters, including cells from both the innate immune compartment, such as border-associated macrophages (BAMs, e.g. *Mrc1* and *Pf4*), classical monocytes (e.g. *Ms4a6c* and *Lgals3*), neutrophils (e.g. *S100a8* and *S100a9*), conventional (cDCs 1 and 2 e.g. *Xcr1* and *Sirpa*, respectively) and monocyte-derived (MoDCs, e.g. *Cd209a*) dendritic cells, natural killer cells (NKs, e.g. *Klrb1c* and *Nkg7)* and ILC2s (e.g. *Il7r* and *Gata3*), as well as the adaptive immune compartment, such as T cells (e.g. *Cd4* and *Cd8b1*) and B cells (e.g. *Ms4a1* and *Cd79a*) (Fig. 2c,d; Extended Data Fig. 2a,b). Notably, we observed a robust increase in several leukocyte subpopulations post-SNI, predominantly within the myeloid compartment, including neutrophils, classical monocytes, BAMs and DCs (cDCs and MoDCs) (Fig. 2e-g; Extended Data Fig. 2c). Additionally, the relative frequencies of ILC2s and T cells increased, NK cells remained unchanged, and B-cell frequency decreased (Fig. 2e,f; Extended Data Fig. 2c). These changes in leukocyte composition in the DRG meninges after SNI were accompanied by significant transcriptional alterations across all cell clusters (Extended Data Fig. 2d).

**Fig. 2.**
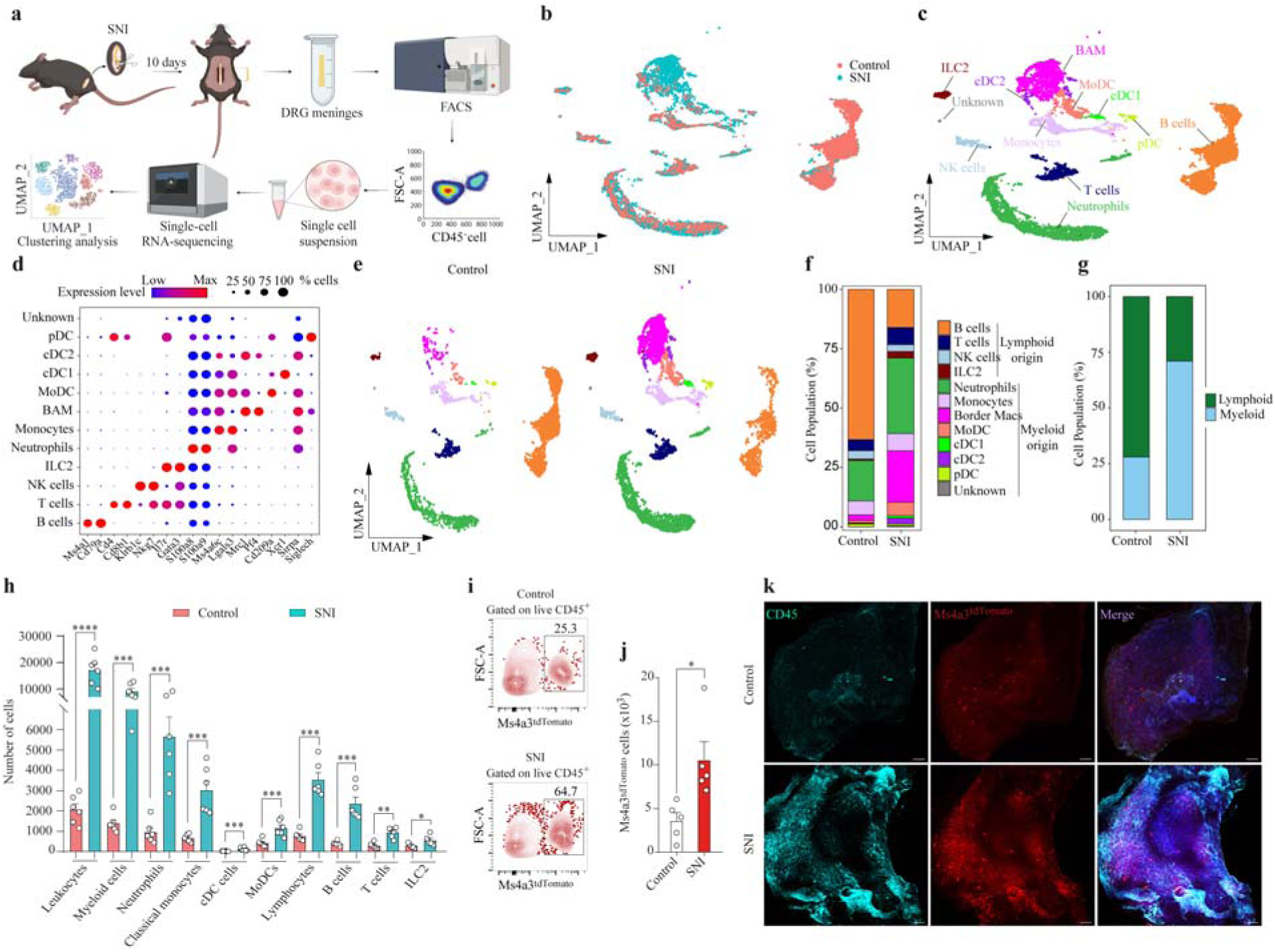
Single-cell transcriptomic atlas of leukocytes in the DRG meninges after peripheral nerve injury. **a.** Schematic illustration of the experimental protocol workflow. C57BL/6J wild-type (WT) mice were subjected to the SNI model (10 mice pooled) or control (20 mice pooled). Ten days later, the DRG meninges were harvested, pooled, and processed for scRNA-seq protocol, dataset collection and analysis. **b.** Uniform Manifold Approximation and Projection (UMAP) representation of the cells present in the DRG meninges comparing control *versus* SNI group. **c.** UMAP representation of the cell population clusters present in the DRG meninges from both control and SNI groups together. Colors are randomly assigned to each cell population. **d.** Dot plot representation of cluster-defining genes for each cell population, where each gene represents the most significant cluster-defining marker for each population. The color and size of each dot represent the average expression and percentage of cells expressing each gene, respectively. **e.** UMAP plot depicting scRNA-seq analysis of leukocytes from the DRG meninges of control or SNI mice. **f.** Stacked bar plot depicting the proportions of cell clusters from data shown in e. Graphs were calculated using Seurat by normalizing the dataset, finding the variable features of the dataset, scaling the data, and reducing the dimensionality. **g.** Stacked bar plot depicting the proportions of myeloid and lymphoid cells from data shown in c. **h.** Absolute number of different cell populations in the DRG meninges from control and SNI samples. *n* = 6 per group. **i.** Flow cytometry gating for *Ms4a3*^tdTomato^ cells in the DRG meninges from *Ms4a3*^tdTomato^ control mice or 10 days after SNI. **j.** Bar graph depicting the number of *Ms4a3*^tdTomato^ cells in the DRG meninges from *Ms4a3*^tdTomato^ control mice or 10 days after SNI. *n* = 5 per group. **k.** Representative images of whole mounts containing L4 DRG and attached meninges stained for CD45^+^ cells (green), *Ms4a3*^tdTomato^ cells (red) and DAPI (blue). Samples were harvested from *Ms4a3*^tdTomato^ mice subjected to SNI (10 days) or control. Scale bars 100 μm. Statistical analysis: unpaired two- sided *t*-tests (h, j). \**P* < 0.05, \*\**P* < 0.01, \*\*\**P* < 0.001, \*\*\*\**P* < 0.0001. n = biologically independent samples from individual mouse tissues. Error bars indicate the mean ± s.e.m. Illustrations were created with BioRender.com (https://biorender.com).

In line with the scRNA-seq results, flow cytometry confirmed and extended these findings, revealing that among leukocyte subsets, myeloid cells exhibited the most pronounced increase in the DRG meninges after SNI, particularly neutrophils and classical (Ly6C^+^) monocytes (Fig. 2h; Extended Data Fig. 2e). The numbers of cDCs and MoDCs were also elevated in the DRG meninges of SNI mice (Fig. 2h; Extended Data Fig. 2e). Within the lymphoid compartment, ILC2s, B cells, CD4^+^ and CD8^+^ T cells were similarly increased (Fig. 2h; Extended Data Fig. 2e, f). SNI induced comparable increases in myeloid and lymphoid cells in male and female DRG meninges (Extended Data Fig. 2g). Furthermore, induction of SNI in *Ms4a3*^tdTomato^ mice, which enable tracking of granulocyte-monocyte progenitor (GMP)- derived cells (monocytes and neutrophils)^25^, revealed an increase in *Ms4a3*^tdTomato^ cells in the DRG meninges after nerve injury (Fig. 2i-k). Neutrophils and macrophages also accumulated in the DRG meninges after CCI (Extended Data Fig. 2h-j). Thus, peripheral nerve injury promotes the accumulation and transcriptional adaptation of innate and adaptive immune cells, particularly myeloid cells, in the DRG meninges.

## Nerve injury induces vertebral emergency myelopoiesis

Meningeal immune cells, particularly myeloid cells, can originate from the skull BM^26,27,28^ and migrate to the meninges through connecting ossified channels in both mice and. Beyond the skull, evidence also suggests that vertebral BM can communicate directly with the spinal cord meninges through similar bone channels^28,32^. To determine whether the myeloid cells accumulating in the DRG meninges after peripheral nerve injury originate from vertebral BM and reach the meninges via vertebral bone channels, we employed several approaches. SEM analysis of the inner surface of the lumbar vertebral arch revealed the presence of microscopic channels (Fig. 3a). High-resolution micro- computed tomography (micro-CT) of the L4 vertebra (Fig. 3b) further demonstrated compact microchannels traversing the bone and connecting the spinal cavity with the vertebral trabecular space (Fig. 3b). These channels were distributed across the dorsal, lateral and ventral regions of the vertebra, with diameters ranging from 8 to 80 μm, and were also visible on the inner surface of the lumbar vertebral arch (Extended Data Fig. 3a). Micro-CT imaging of human vertebral bone similarly revealed channels connecting the inner vertebral surface with the underlying marrow cavities (Fig. 3c).

**Fig. 3.**
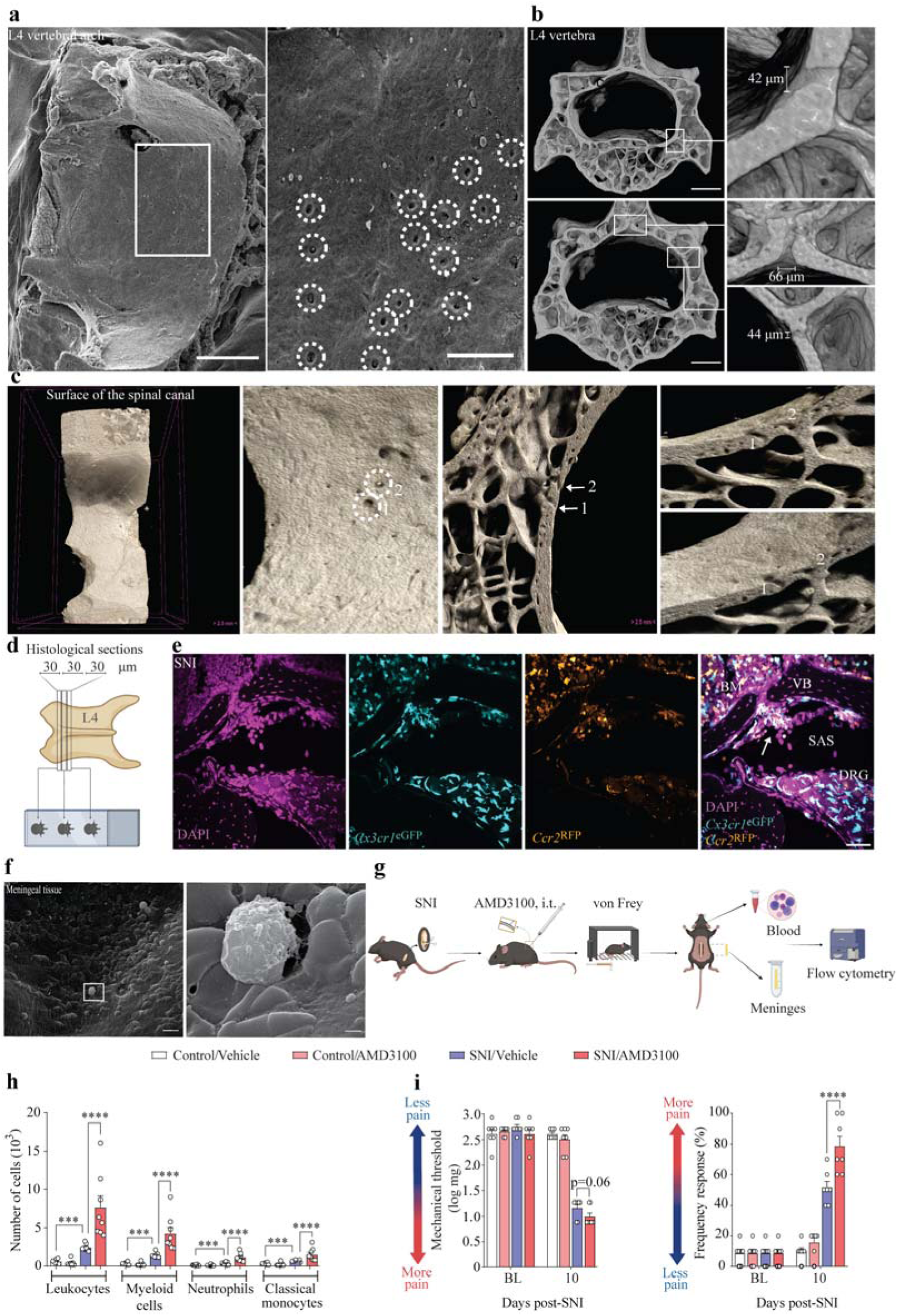
Peripheral nerve injury triggers myeloid cell migration towards DRG meninges through vertebral ossified channels. **a.** SEM image of the inner surface of the L4 vertebral arch. Ossified channels openings (dash circles) on the bone surface (right panel). Scale bars, 500 μm (left) and 100 μm (right). **b.** Representative maximum intensity projection of the transversal view of the lumbar vertebra (L4) imaged by μ-CT (left panels). High magnification showing ossified channels in the L4 vertebral arch connecting marrow cavity and spinal canal (right) Scale bars, 500 µm (left) and 50 μm (right). **c.** Digital section in the sagittal plane of a representative human lumbar vertebra imaged by μ-CT. The dashed circles (1 and 2) in the left panel indicate the opening of the ossified channel crossing the cortical bone. Arrows (right panels) show the channels connecting marrow cavity and spinal canal. Scale bars, 2.5 mm. **d.** Schematic illustration of the protocol to prepare histological specimens from the whole lumbar spinal column. Three consecutive sections (30 μm each) were sliced in the transversal plane and prepared for confocal imaging. **e.** Representative confocal image showing *Ccr2*^RFP^ and *Cx3cr1*^eGFP^ positive leukocytes leaving (marked by arrow) the vertebral BM toward subarachnoid space (SAS) through vertebral bone (VB) ossified channel near the L4 DRG. Sample collection was performed 10 days after SNI surgery in *Ccr2*^RFP^ *- Cx3cr1*^eGFP^ dual reporter mice. Scale bar, 50 μm. **f.** Scanning electron microscopy image of the outer surface of mouse dorsal dura mater in the DRG meninges. The insertion frame and high magnification (right panel) are showing leukocyte crossing a dura mater membrane pore. Sample collection was performed 10 days after SNI. Scale bar, 10 μm (left) and 1 μm (right). **g.** Schematic illustration of the experimental protocol workflow. C57BL/6J wild-type (WT) mice were subjected to the SNI model. Seven and nine days later, they received an i.t. injection of AMD3100 (5 μg/site), or vehicle followed by pain behaviors analyses (von Frey test) and isolation of DRG meninges and blood for flow cytometry analyses. **h.** Absolute number of different cell populations in the DRG meninges from control and SNI samples that receive vehicle or AMD3100 (5 μg/site, i.t.) treatments. *n* = 8 per group. **i.** Analyses of mechanical pain hypersensitivity by *von Frey* tests in control and SNI mice that receive vehicle or AMD3100 (5 μg/site, i.t.) treatments. *n* = 8 per group. Statistical analysis: one-way ANOVA with Tukey post-tests (h, i). \*\*\**P* < 0.001, \*\*\*\**P* < 0.0001, ns = non-significant. n = biologically independent samples from individual mice or mouse tissues. Error bars indicate the mean ± s.e.m. Illustrations were created with BioRender.com (https://biorender.com).

To visualize the migration of myeloid cells from the vertebral BM to the DRG meninges after SNI, we used *Cx3cr1*^eGFP^-*Ccr2*^RFP^ dual-reporter mice. In whole-spinal column preparation containing the L4 DRG, vertebral bone, BM, and subarachnoid space (Fig 3d), we observed classical monocytes leaving the vertebral BM and accessing the subarachnoid space (Fig 3e; Extended Data Fig. 3b), where they clustered in the L4 DRG meninges (Extended Data Fig. 3c). SEM images acquired from the outer dural surface also revealed cells crossing the dura mater (Fig 3f). Finally, to confirm the direct recruitment of myeloid cells to the DRG meninges from the vertebral BM through vertebral bone channels after SNI, we intrathecally (i.t.) injected a low dose of AMD3100, a CXCR4 antagonist (Fig. 3g), which promotes the release of myeloid cells from the BM by blocking the CXCL12 (SDF-1)-CXCR4 retention mechanism^33^. This strategy has previously been used to demonstrate the mobilization of skull BM cells into the brain meninges^30,34^. The injection of AMD3100 in SNI mice enhanced both the accumulation of myeloid cells in the DRG meninges and pain hypersensitivity (Fig. 3h,i). Notably, AMD3100 did not alter blood leukocyte counts (Extended Data Fig. 3d,e), ruling out a possible systemic effect. These findings support the conclusion that myeloid cells can egress from the vertebral BM to the DRG meninges through vertebral bone channels.

In various CNS pathologies, including Alzheimer’s disease, multiple sclerosis, stroke, and meningitis, skull BM emergency myelopoiesis can supply the brain meninges with *de novo* myeloid cells^28,29^,

To determine whether vertebral myelopoiesis is similarly induced after peripheral nerve injury, we performed flow cytometry on lumbar vertebral BM cells, using tibial BM as a control for potential systemic changes in myelopoiesis (Fig. 4a). Vertebral BM niches displayed an increase in both the number and proliferation of common myeloid progenitors (CMPs) and GMPs after SNI compared with control mice, indicating the induction of emergency myelopoiesis (Fig. 4b-d; Extended Data Fig. 4a). Moreover, the numbers of neutrophils and classical monocytes were markedly increased in the vertebral BM after nerve injury (Fig. 4e; Extended Data Fig. 4b). We also observed a reduction in the expression of *Cxcl12* (encoding CXCL12/SDF-1), providing evidence for decreased retention of myeloid cells within the vertebral BM after SNI (Fig. 4f). Importantly, no alterations in cellular composition were observed in the tibial BM (Extended Data Fig. 4c-g), indicating no major systemic changes in myelopoiesis after SNI.

**Fig. 4.**
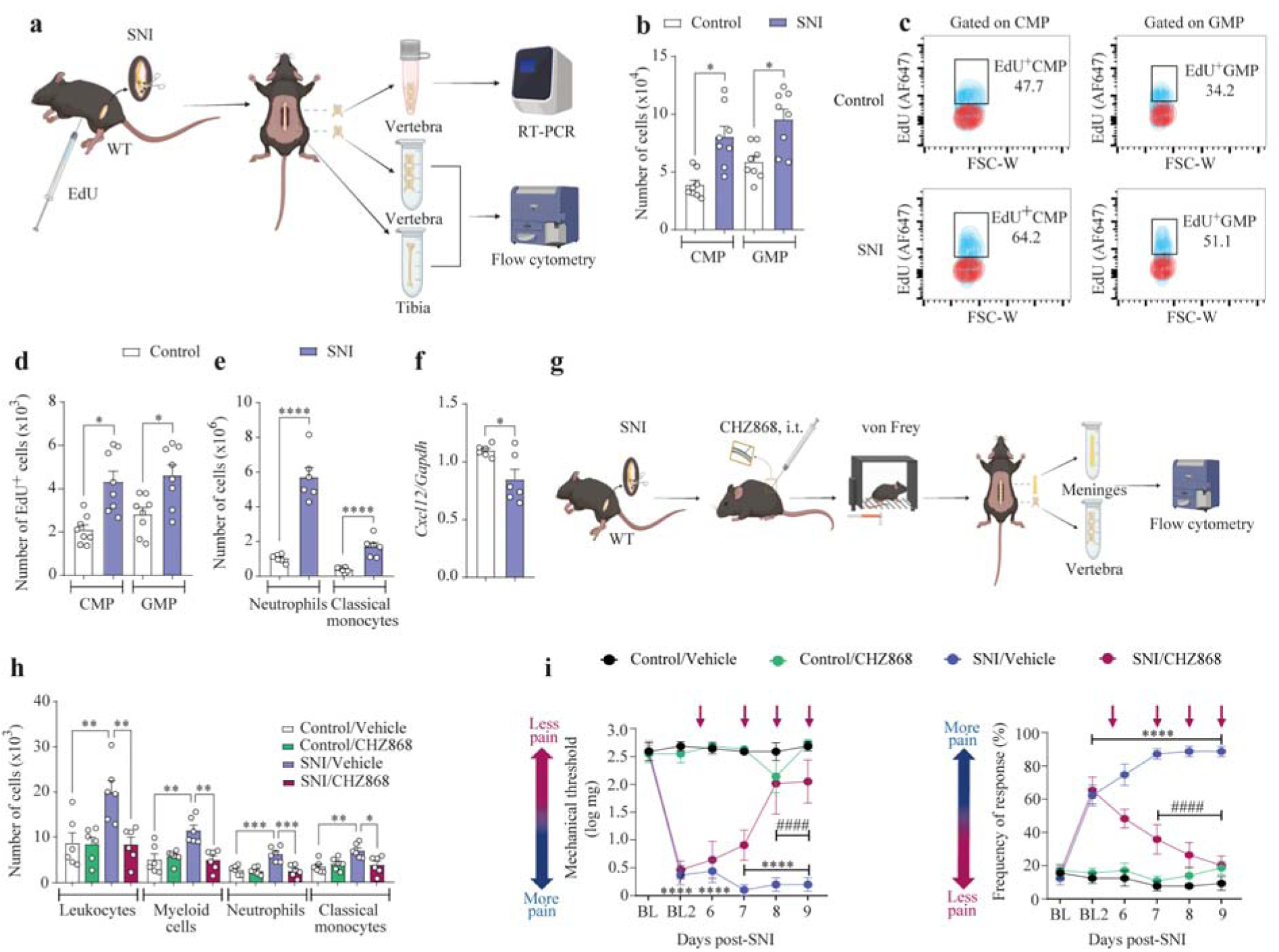
Peripheral nerve injury stimulates vertebral BM emergency myelopoiesis- dependent neuropathic pain. **a.** Schematic illustration of the experimental protocol workflow. C57BL/6J wild-type (WT) mice were subjected to the SNI model. Ten days later, the vertebral and tibial BM was harvested, and processed for flow cytometry or RT-PCR. Tissues from control mice were used as controls. **b.** Absolute number of common myeloid progenitors (CMP) and granulocyte-monocyte progenitor (GMP) in the vertebral BM from control mice or 10 days after SNI. *n* = 8 mice per group. **c.** Representative flow cytometry gating plots showing EdU^+^CMP and EdU^+^GMP in the vertebral BM from control or 10 days after SNI. **d.** Absolute number of EdU^+^CMP and EdU^+^GMP in the vertebral BM from control or 10 days after SNI. *n* = 8 mice per group. **e.** Absolute number of neutrophils and classical monocytes in the vertebral BM from control mice or 10 days after SNI. *n* = 6 per group. **f.** qPCR analysis of relative expression of C*xcl12 in* the vertebral BM from control mice or 10 days after SNI. *n* = 6 mice per group. **g.** Schematic illustration of the experimental protocol workflow. C57BL/6J wild-type (WT) mice were subjected to the SNI model. Six to nine days later, they received an i.t. injection of CHZ868 (20 μg/site), or vehicle followed by pain behaviors analyses (von Frey filament test) and isolation of DRG meninges and vertebral BM for flow cytometry analyses. **h.** Absolute number of different cell populations in the DRG meninges from control and SNI samples that received vehicle or CHZ868 (20 μg/site) treatments. *n* = 6 per group. **i.** Analyses of mechanical pain hypersensitivity by *von Frey* tests in control and SNI mice that received vehicle or CHZ868 (20 μg/site) treatments. BL = baseline; BL2 = mechanical pain threshold 6 days post-SNI and before treatment. *n* = 6 per group. Statistical analysis: two-way ANOVA with Bonferroni post-test (i), one-way ANOVA with Tukey post-tests (h), unpaired two-sided *t*-tests (b, d, e, f). \**P* < 0.05, \*\*\**P* < 0.001, \*\*\*\**P* < 0.0001, ^####^*P < 0.001*. *n* = biologically independent samples from individual mice or mouse tissues. Error bars indicate the mean ± s.e.m. Illustrations were created with BioRender.com (https://biorender.com).

To assess whether myeloid cell accumulation in the DRG meninges depends on vertebral BM emergency myelopoiesis, we locally inhibited emergency myelopoiesis in SNI mice using intrathecal injection of CHZ868 (Fig. 4g)^35^, a JAK2-selective inhibitor belonging to a new generation of drugs approved for myeloproliferative neoplasms^36^. CHZ868 treatment blocked the SNI-induced expansion of CMPs and GMPs, as well as their derived myeloid cells, in the vertebral BM (Extended Data Fig. 4h,i). Inhibition of emergency myelopoiesis with CHZ868 also reduced the SNI-induced accumulation of myeloid cells in the DRG meninges and the associated pain hypersensitivity (Fig. 4h,i). Altogether, these findings demonstrate that peripheral nerve injury triggers BM emergency myelopoiesis, which supplies myeloid cells to the DRG meninges to promote pain hypersensitivity.

## GM-CSF stimulates vertebral emergency myelopoiesis

Induction of skull BM emergency myelopoiesis has been observed in several pathological conditions^37^, however, the specific signal triggering this process remains unknown. To identify potential molecular signal inducing vertebral BM emergency myelopoiesis after peripheral nerve injury, we first performed pathway enrichment analysis using gene ontology on all clusters of DRG meningeal leukocytes identified in the scRNA-seq (Extended Data Fig. 5a). Only ILC2s showed significant enrichment for pathways related to the regulation of hematopoiesis (Extended Data Fig. 5b). Interestingly, imaging of whole-mount DRG (L4) after SNI revealed that ILC2s were located exclusively in the surrounding meninges (Extended Data Fig. 5c). To explore the translational relevance of these findings, we reanalyzed^38^ spatial RNA-sequencing data from DRGs of organ donors who died with a history of painful diabetic neuropathy (PDN), the most common form of neuropathic pain in humans^39^. We found an increased number of barcodes with ILC2 markers in PDN DRGs compared to controls, and these barcodes were located either at the edge of the DRG or in areas associated with the DRG meninges, consistent with our mouse findings (Extended Data Fig. 5d).

To identify ILC2-derived signals that could affect myelopoiesis, we used the CellChat algorithm to infer potential cell-cell interactions. Among the significant interactions identified between ILC2s and myeloid cells, one associated with emergency myelopoiesis and appearing specifically after SNI was the *Csf2*–*Csf2ra/Csf2rb* axis (Extended Data Fig. 6a). *Csf2* encodes GM-CSF, an emergency myelopoietic growth factor and pro-inflammatory cytokine that promotes the activation and polarization of myeloid cells, including neutrophils and monocytes/macrophages^40,41^, whereas *Csf2ra* and *Csf2rb* encode its cellular receptor subunits, CSF2RA and CSF2RB, which dimerize after GM-CSF binding^40,41^. This communication appeared highly selective, originating from ILC2s toward myeloid cells but not lymphoid cells (Extended Data Fig. 6b). Notably, *Csf2* was one of the most enriched transcripts in ILC2s compared to other leukocyte subpopulations in the DRG meninges (Extended Data Fig. 6c), being almost exclusively expressed in this cluster, whereas *Csf2ra* and *Csf2rb* were detected only in myeloid cells (Fig. 5a).

**Fig. 5.**
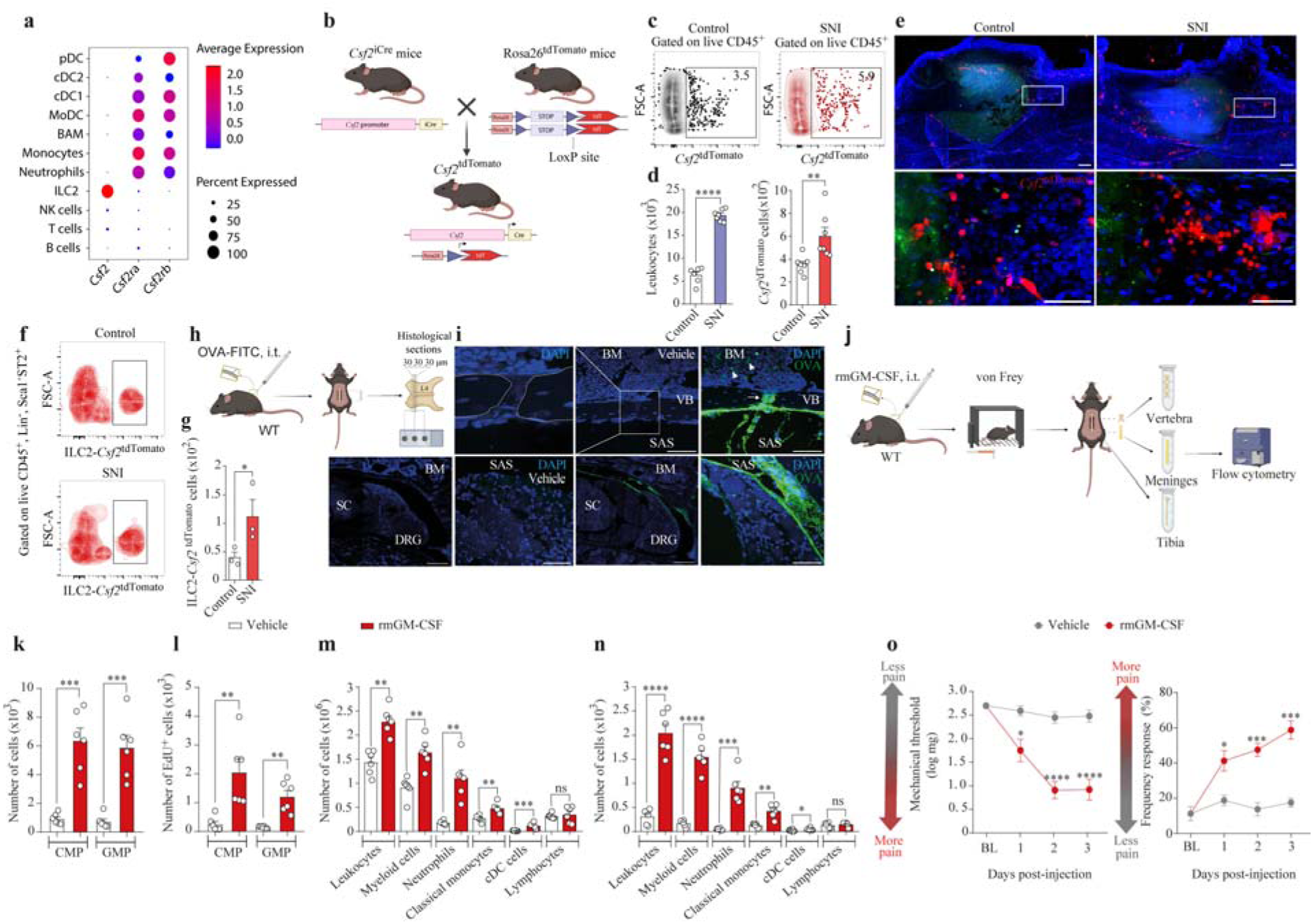
Meningeal GM-CSF instructs vertebral BM emergency myelopoiesis-dependent pain hypersensitivity. **a.** Dot plot of *Csf2*, *Csf2ra*, and *Csf2rb* expression across immune cell types in DRG meninges. **b.** Schematic illustration of the generation of *Csf2*^tdTomato^ experimental mouse model by breeding *Csf2*^iCre^ mice with *Rosa26*^tdTomato^ mice. **c.** Flow cytometry gating for *Csf2*^tdTomato^ cells in the DRG meninges from control mice or 10 days after SNI. **d.** Absolute number of leukocytes (CD45^+^ cells, left panel) and *Csf2*^tdTomato^ cells (right panel) in DRG meningeal tissue from control mice or 10 days after SNI. *n* = 7 per group. **e.** Representative images of meningeal DRG’s whole mount from control *Csf2*^tdTomato^ (left) and SNI *Csf2*^tdTomato^ (right) mice. DAPI (blue) for nucleus staining. Scale bars (from top to bottom), 200 µm and 50 μm. **f.** Flow cytometry gating for ILC2-*Csf2*^tdTomato^ cells in the DRG meninges from control mice or 10 days after SNI. **g.** Absolute number of ILC2- *Csf2*^tdTomato^ cells in the DRG meninges from control mice or 10 days after SNI. *n* = 3 per group (pooled from 3 mice). **h.** Schematic illustration of the experimental protocol workflow. C57BL/6J wild-type (WT) mice received i.t. injection of OVA-FITC or vehicle. Twenty-four hours later, lumbar spinal column tissue was harvested and processed for confocal analyses. **i.** Representative confocal image showing OVA-FITC staining of meningeal tissue after i.t. injection. Arrowheads highlight OVA-FITC accumulation in the vertebral bone channel. Arrows denote OVA-FITC^+^ cells within the vertebral bone marrow (BM). Scale bars, 50 μm. **j.** Schematic illustration of the experimental protocol workflow. C57BL/6J wild-type (WT) mice received i.t. injection of recombinant murine (rm) GM-CSF (100 ng/site/injection) or vehicle for three days, twice a day, followed by pain behaviors analyses (von Frey filament tests) and the isolation of DRG meninges, vertebral BM or tibia BM for flow cytometry analyses. **k.** Absolute number of common myeloid progenitors (CMP) and granulocyte- monocyte progenitor (GMP) in the vertebral BM from control mice that received vehicle or rmGM-CSF (100 ng/site/injection). *n* = 6 per group. **l.** Absolute number of EdU^+^ CMP and EdU^+^ GMP in the vertebral BM from mice that received vehicle or rmGM-CSF. *n* = 6 per group. **m.** Absolute number of different cell populations in the vertebral BM tissue from mice that received vehicle or rmGM-CSF. *n* = 6 per group. **n.** Absolute number of different cell populations in the DRG meninges from mice that received vehicle or rmGM-CSF. *n* = 6 per group. **o.** Mechanical pain hypersensitivity before (baseline, BL) and 6 hours after each rm GM-CSF injection. *n* = 6 per group. Statistical analysis: two-way ANOVA with Bonferroni Bonferroni (o), unpaired two-sided *t*-tests (d, g, k-n). \**P* < 0.05, \*\*\**P* < 0.001, \*\*\*\**P* < 0.0001, ns = non-significant. n = biologically independent samples from individual mice or mouse tissues. Error bars indicate the mean ± s.e.m. Illustrations were created with BioRender.com (https://biorender.com).

To confirm the presence of *Csf2*^+^ cells in the DRG meninges, we crossed mice expressing iCre recombinase under the *Csf2* promoter (*Csf2*^iCre^) with the *Rosa26*^tdTomato^ reporter line (referred to as *Csf2*^tdTomato^, Fig. 5b)^42^. The number of *Csf2*^tdTomato^ cells, gated on CD45^+^, increased after SNI in the DRG meninges (Fig. 5c,d), whereas their number was minimal in non-leukocytic compartment (less than 0.1% of CD45^-^ cells) and did not change after SNI (Extended Data Fig. 6d). In whole-mount preparation, *Csf2*^tdTomato^ cells were predominantly localized within the meninges surrounding injured DRGs (Fig. 5e) and were absent in the parenchyma of DRG or spinal cord (Extended Data Fig. 6e,f). Consistent with the scRNA-seq data, the number of ILC2s expressing *Csf2*^tdTomato^ increased in the DRG meninges after SNI (Fig. 5f,g). The expression of *Csf2* in the DRG meninges of *Rag1*^-/-^ mice (lacking T and B cells) was similar to controls (Extended Data Fig. 6g), indicating that *Csf2* is not produced by T and B cells in DRG meninges. Based on these findings and previous studies showing that ILC2s have a high capacity to produce GM-CSF^43,44^, we hypothesized that GM-CSF is a soluble factor released after peripheral nerve injury, primarily from meningeal ILC2s, that instructs emergency myelopoiesis in the vertebral BM.

To assess whether GM-CSF can reach the vertebral BM, we first injected fluorescent ovalbumin (OVA-FITC, ∼40 kDa) intrathecally and examined its efflux to the vertebral BM, as previously reported^30^ (Fig. 5h). Imaging of spinal column preparation revealed tracer distribution along the meningeal layers, minimal signal in the DRG parenchyma, and no staining in the spinal cord parenchyma (Fig. 5i). Importantly, the tracer was present within the ossified channels in the vertebral bone and within the BM niche (Fig. 5i), indicating that proteins released from the meninges can reach the vertebral BM. In agreement, intrathecal injection of recombinant GM-CSF promoted vertebral BM emergency myelopoiesis, as evidenced by the expansion of CMPs, GMPs and myeloid cells (Fig. 5j-m). No changes were observed in the tibia BM after the intrathecal injection of GM-CSF, demonstrating a local effect in the vertebral BM compartment (Extended Data Fig. 6h). Consistent with the stimulation of emergency myelopoiesis, intrathecal GM-CSF also increased the number of leukocytes, particularly myeloid but not lymphoid cells, in the DRG meninges (Fig. 5n) and induced pain hypersensitivity (Fig. 5o). Notably, the GM-CSF–induced increase in leukocyte accumulation in the DRG meninges and pain hypersensitivity was abolished by inhibiting vertebral emergency myelopoiesis (using JAK2 inhibitor, Extended Data Fig. 6i–k). These findings demonstrate that GM-CSF induces leukocyte accumulation in the DRG meninges and promotes pain hypersensitivity through the induction of vertebral BM emergency myelopoiesis.

To further study the role of the GM-CSF–CSF2R axis in mediating emergency myelopoiesis after peripheral nerve injury and the resulting neuropathic pain, we generated a knock-in reporter mouse expressing fluorescent ZsGreen under the control of the *Csf2ra* promoter (referred to as *Csf2ra*^ZsGreen^; the targeting strategy and validation are shown in Extended Data Fig. 7a-d) and mice lacking CSF2RA (referred to as *Csf2ra^-/-^;* the targeting strategy and validation are shown in Extended Data Fig. 7a-c,e). Following SNI, *Csf2ra*^ZsGreen^ cells expanded in the vertebral BM (Fig. 6a-c; Extended Data Fig. 8a,b) and accumulated in the DRG meninges. The majority of *Csf2ra*^ZsGreen^ cells in both sites were myeloid cells (>95%), consisting primarily of neutrophils (55%) and classical monocytes (38%) (Extended Data Fig. 8c,d). *Csf2ra*^ZsGreen^ cells were absent in the DRG (Extended Data Fig. 8a) and spinal cord (Extended Data Fig. 9b) parenchyma.

**Fig. 6.**
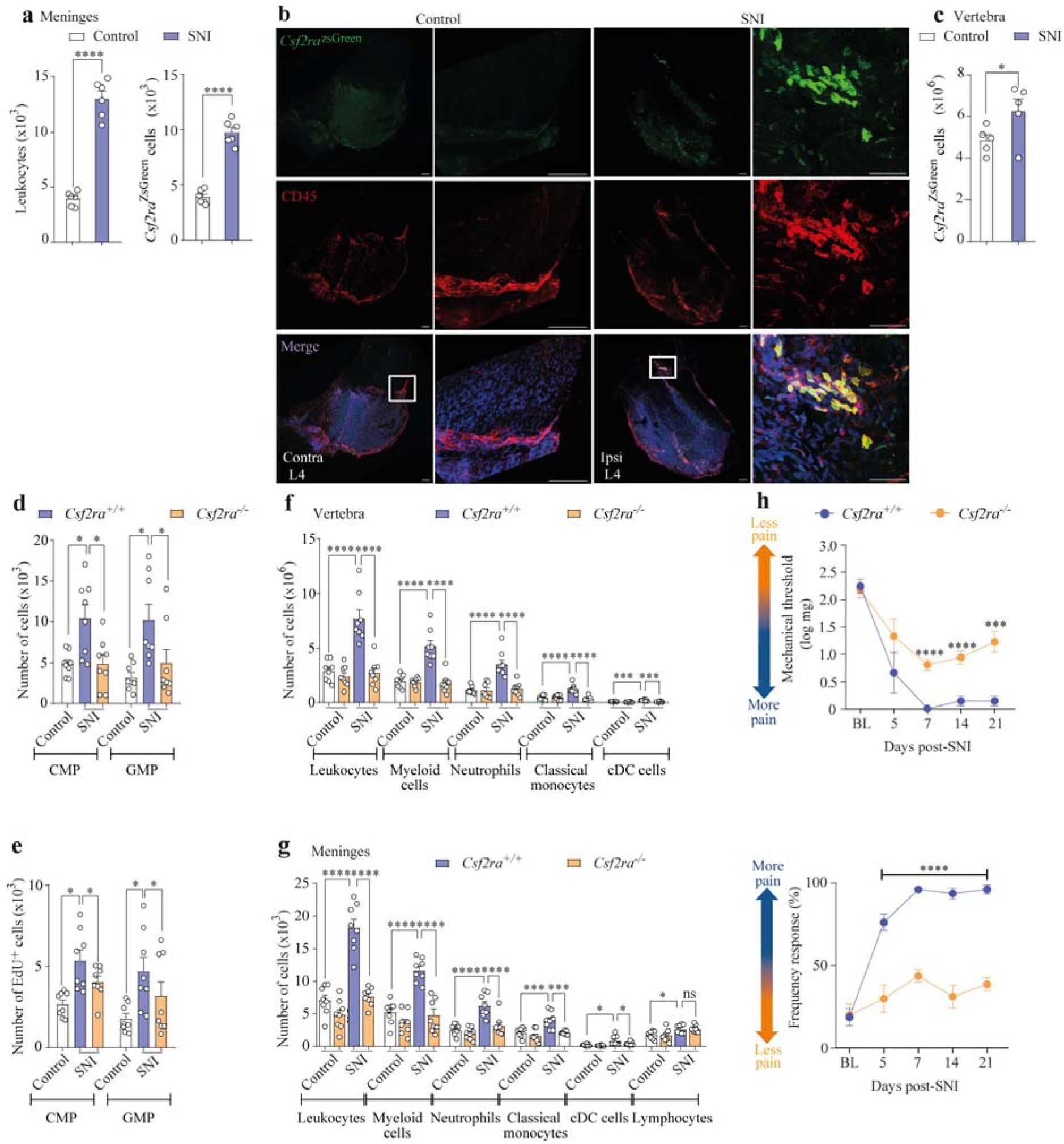
GM-CSF-CSFR signaling mediates neuropathic pain through induction of emergency myelopoiesis. **a.** Absolute number of leukocytes (CD45^+^ cells, left panel) or *Csf2ra*^ZsGreen^ cells in the DRG meninges from *Csf2ra*^ZsGreen^ mice subjected to SNI (10 days) or control. n = 6 per group. **b.** Representative images of meningeal DRG whole mounts stained for CD45^+^ cells (red), *Csf2ra*^ZsGreen^ cells (green), and DAPI (blue). Samples were harvested from *Csf2ra*^ZsGreen^ mice subjected to SNI (10 days) or control. Scale bars (from left to right), 100 μm and 50 μm. **c.** Absolute number of *Csf2ra*^ZsGreen^ cells in the vertebral BM from *Csf2ra*^ZsGreen^ mice subjected to SNI (10 days) or control. n = 5 per group. **d.** Absolute number of common myeloid progenitors (CMPs) and granulocyte-monocyte progenitor (GMPs) in the vertebral BM from *Csf2ra*^+/+^ or *Csf2ra*^-/-^ control mice or 10 days after SNI. *n* = 8 per group. **e.** Absolute number of EdU^+^ CMP and EdU^+^ GMP in the vertebral BM from *Csf2ra*^+/+^ or *Csf2ra*^- /-^ control mice or 10 days after SNI. *n* = 8 per group. **f.** Absolute number of different cell populations in the vertebral BM from *Csf2ra*^+/+^ or *Csf2ra*^-/-^ control mice or 10 days after SNI. *n* = 8 per group. **g.** Absolute number of different cell populations in the DRG meninges from *Csf2ra*^+/+^ or *Csf2ra*^-/-^ control mice or 10 days after SNI. *n* = 8 per group. **h.** Time course of mechanical pain hypersensitivity in *Csf2ra*^+/+^ or *Csf2ra*^-/-^ mice before (baseline, BL) and after SNI. *n* = 8 per group. Statistical analysis: two-way ANOVA with Bonferroni post-test (h); one-way ANOVA with Tukey post-tests (d-g), unpaired two-sided *t*-tests (a, c). \**P* < 0.05, \*\**P* < 0.01, \*\*\**P* < 0.001, \*\*\*\**P* < 0.0001. ns = non-significant. *n* = biologically independent samples from individual mice or mouse tissues. Error bars indicate the mean ± s.e.m.

In *Csf2ra^-/-^* mice, intrathecal GM-CSF failed to stimulate vertebral BM emergency myelopoiesis and did not induce the accumulation of myeloid cells in the DRG meninges (Extended Data Fig. 10a-e). *Csf2ra^-/-^*mice exhibited no changes in basal pain sensitivity or motor function. Importantly, intrathecal GM-CSF induced pain hypersensitivity in control animals but not in *Csf2ra^-/-^* mice (Extended Data Fig. 10f,g). These findings indicate that GM- CSF drives vertebral myelopoiesis and pain hypersensitivity through its cognate receptor, CSF2R.

In the SNI model of neuropathic pain, *Csf2ra*^-/-^ mice exhibited reduced vertebral BM emergency myelopoiesis, decreased accumulation of myeloid cells (neutrophils and classical monocytes) in the DRG meninges, and attenuated pain hypersensitivity (Fig. 6d-h). Consistent with the lack of *Csf2ra* expression in the DRG parenchyma, no differences were observed in the responses of DRG parenchymal macrophages in *Csf2ra* ^/^ mice after SNI (Extended Data Fig. 11a,b). In addition, GM-CSF did not alter the excitability of cultured sensory neurons, as assessed by patch-clamp and microelectrode array (MEA) recordings (Extended Data Fig. 12a,b), excluding a direct effect on primary sensory neurons and consistent with previous findings^45^. Similar to *Csf2ra^-/-^*mice, GM-CSF knockout animals (*Csf2* ^/^) also exhibited reduced vertebral BM emergency myelopoiesis, diminished accumulation of myeloid cells in the DRG meninges, and attenuated pain hypersensitivity after SNI (Extended Data Fig. 13a–e), without alterations in baseline nociception (Extended Data Fig. 13f). Altogether, these results indicate that meningeal GM-CSF, acting through GM-CSFR signaling, induces vertebral BM emergency myelopoiesis, leading to the accumulation of myeloid cells in the DRG meninges and the development of neuropathic pain.

## Discussion

Here, we reveal a neuroimmune circuit that links the DRG meninges–vertebral BM axis to pathological pain. Following peripheral nerve injury, the vertebral BM undergoes emergency myelopoiesis, supplying myeloid cells to the DRG meninges via ossified vertebral bone channels. Mechanistically, meningeal GM-CSF instructs vertebral BM emergency myelopoiesis, thereby promoting the development of neuropathic pain (Extended Data Fig. 14).

After peripheral nerve injury, cells within sensory ganglia (DRG or trigeminal ganglia) produce various pro-nociceptive mediators, such as cytokines and chemokines, creating a pathological environment that promotes hyperexcitability of primary sensory neurons^8^. Whereas parenchymal immune interactions have been well characterized^46,47,48^, the immune landscape of the DRG meninges is only beginning to be understood. We found that immune cells of both innate and adaptive lineages accumulate in the meninges covering injured DRGs. The selective accumulation of immune cells in the meninges of the DRGs associated with the injured sciatic nerve suggests local release of factors from these DRGs that guide leukocyte recruitment. Chemokines such as CXCL1 and CCL2, which are upregulated in the DRG after peripheral nerve injury and can attract neutrophils and monocytes, respectively, are plausible candidates^48,49,50,51,52^.

CNS damage and disease induce the recruitment of myeloid cells into the brain meninges, accompanied by skull BM emergency myelopoiesis and their migration through skull ossified channels^26,37,52^. We now provide direct evidence that vertebral BM emergency myelopoiesis supplies myeloid cells to the DRG meninges via vertebral bone channels. Moreover, we show that this process functionally contributes to neuropathic pain. The accumulated meningeal myeloid cells could enhance sensory neuron excitability and pain signaling through numerous mechanisms, including the release of pro-nociceptive mediators such as TNF, IL-1β, chemokines, lipid mediators, and reactive oxygen species^8^.

Importantly, we identified GM-CSF, produced by meningeal ILC2s, as a key inducer of vertebral emergency myelopoiesis after peripheral nerve injury. The ability of meningeal ILC2–derived GM-CSF to elicit a hematopoietic response reveals a previously unappreciated coupling between barrier-resident lymphoid cells and vertebral BM function. This may represent a conserved neuroimmune axis linking nervous tissue inflammation or disease to local myelopoietic adaptation across CNS pathologies^28,29,52,53^.

The translational relevance of our findings is supported by several lines of evidence in humans. The presence of ossified vertebral bone channels in humans suggests that direct immune cell trafficking between vertebral BM and meninges may occur in patients. Clinical trials have shown that anti-GM-CSF antibodies rapidly alleviate pain in rheumatoid arthritis, whereas GM-CSF administration to promote bone marrow recovery frequently exacerbates chemotherapy-induced neuropathic pain^59,60,61,62^ . A recent study reported an increase in TSPO signal in the skull BM of chronic pain patients, indicating elevated immune activity^63^. Finally, we observed an increase in ILC2s in human DRGs from individuals with a history of PDN, where ILC2-positive regions were localized to the outer DRG layers, consistent with an expansion of meningeal ILC2s in humans with neuropathic pain. These observations are consistent with a model in which GM-CSF (primarily from ILC2s)–driven emergency myelopoiesis enhances the recruitment and activation of meningeal myeloid cells, thereby sustaining neuroinflammatory circuits that amplify nociceptive signaling. They also support the existence of a conserved BM–meningeal axis that links hematopoietic activity to chronic pain pathophysiology in humans.

In summary, we identified a previously unrecognized meningeal–vertebral BM axis that shapes the neuroimmune landscape after peripheral nerve injury. GM-CSF released from the meninges instructs local vertebral BM emergency myelopoiesis, leading to myeloid cell accumulation in the DRG meninges and promoting neuropathic pain. This discovery adds a new dimension to the understanding of meningeal immunity and uncovers potential therapeutic targets for chronic pain through the modulation of meningeal cells and their mediators.

## Methods

### Experimental Animals

All mice used in this study were kept on a C57BL/6J background and were maintained under specific pathogen-free (SPF) conditions in the Center for Transgenic Mice of the Ribeirao Preto Medical School (CCCE-FMRP). In general, littermates were (8-12 weeks old) paired in sex, age and weight for each experiment. The following strains were used: *Wild type* (WT, C57BL6/J mice, Jackson Laboratory, Strain: 000664), *Csf2* (*Csf2^-/-^*mice, Jackson Laboratory, B6.129S-*Csf2tm1Mlg*/J, Strain 026812)^62^, and *Rag1* knockout mice (*Rag1^-/-^* mice, Jackson Laboratory, B6.129S7-Rag1tm1Mom/J, Strain 002216)^63^ *Rosa26*^tdTomato^ (B6.Cg-Gt(ROSA)26Sortm14(CAG-tdTomato)Hze/J Jackson Laboratory, Strain :007914)^64^, *Scn10a*^CRE^ mice^65^ (B6.129(Cg)-Scn10atm2(cre)Jwo/TjpJ, Jackson Laboratory, Strain: 036564) and *Il5^i^*^CRE^ mice^66^ (B6(C)-Il5tm1.1(icre)Lky/J, Jackson Laboratory, Strain: 030926). *Ccr2*^RFP^*-Cx3cr1*^eGFP^ dual-reporter mice is a double mutant strain produced by crossing *Ccr2*^RFP^^67^ (B6.129(Cg)-Ccr2tm2.1Ifc/J, Jackson Laboratory, Strain: 017586) and *Cx3cr1*^eGFP^^68^ (B6.129P-Cx3cr1tm1Litt/J, Jackson Laboratory, Strain: 005582). The *Scn10a*^CRE^, *Csf2*^iCRE42^, *Ms4a3*^CRE25^ and *Il5^i^*^CRE^ knockin mice were bred with *Rosa26*^tdTomato^ mice to generate the *Scn10a*^tdTomato^, *Csf2*^tdTomato^, *Ms4a3*^tdTomato^ and *Il5*^tdTomato^ reporter mice strains. *Csf2ra* knockin (*Csf2ra*^ZsGreen^) and *Csf2ra* knockout mice (*Csf2ra*^-/-^) were designed and generated by genOway for GSK as described below. Both male and female age-matched mice from 8 to 14 weeks of age were used for all experiments in this study. All experimental procedures were approved by University of Sao Paulo Animal Ethical Committee (protocol 142/2020 and 1270/2023) and carried out in accordance with the European Directive 2010/63/EU, UK Home Office Animal Procedures Act (1986), ARRIVE guidelines and the GSK Policy on the Care, Welfare and Treatment of Laboratory Animals.

### Generation of the *Csf2ra* transgenic mouse strains

*Csf2ra^ZsGreen^* and *Csf2ra*^-/-^ mice were designed and generated by genOway (Lyon, France) for GSK. *Csf2ra^ZsGreen^*mice enable *Csf2ra* expression monitoring and its spatio-temporal ablation. They have an IRES-ZsGreen transgenic construct inserted in exon 12 (NM_009970 *Csf2ra* transcript), downstream of the STOP codon (Chr19:61,213,427; GRCm39 Assembly), and two *lox*P sites, in the 5’ (Chr19:61,217,850) and 3’ (Chr19:61,211,809) regions of *Csf2ra*. *Csf2ra*^-/-^ mice have a region between these two *loxP* sites (flanking the entire *Csf2ra* gene) deleted.

The generation of these mice is described in Extended Fig. 6a. Briefly, it included several steps. First, a 5’ *loxP* site was inserted in the *Csf2ra* locus in C57BL/6J embryonic stem (ES) cells (Extended Fig. 6a). Second, a targeting vector including IRES-ZsGreen and a 3’ *loxP* site was inserted downstream of the STOP codon of *Csf2ra* in these modified ES cells using homologous recombination (Extended Fig. 6a), followed by blastocyst injection and generation of chimeras. And finally, the C*sf2ra^ZsGreen^* allele was generated by the *in vivo* deletion of the *loxP-flanked* Neo cassette using Dre-recombinase (Extended Fig. 6a), and the C*sf2ra^-/-^* allele was generated by the *in vivo* deletion of the *loxP*-flanked region using Cre- recombinase (Extended Fig. 6a).

PCR genotyping was conducted using two pairs of primers. Forward primer 1 (ACGTATTGTTCCAGGTCACTTCTGTTTCTCC) and reverse primer 1 (GTTGACATACTTCAAATGGTAAGAACAGTTAGCCA) amplified an 849 bp product for the wild-type allele and a 964 bp product for the C*sf2ra^ZsGreen^* allele. Forward primer 2 (TGCAATTCCCCACAATACCATATTGTCAG) and reverse primer 2 (TACACCTGCACCAGACTCTTTCCTTGTACC) amplified a 180 bp product for the C*sf2ra^-/+^* allele.

### Neuropathic pain models

#### Spared nerve injury (SNI)

A model of neuropathic pain caused by peripheral nerve injury in mice was induced as previously described^23^. Briefly, under isoflurane (2%) anesthesia the skin on the lateral surface of the thigh was incised and a section made directly through the biceps femoris muscle exposing the sciatic nerve and its three terminal branches: the sural, common peroneal and tibial nerves. The SNI comprises an axotomy and ligation of the tibial and common peroneal nerves leaving the remaining sural nerve intact. The common peroneal and the tibial nerves were tightly ligated with 5.0 silk and sectioned distal to the ligation, removing 2 ± 4 mm of the distal nerve stump. Sham-operated or naive animals were used as a control.

#### Chronic constriction injury (CCI)

Neuropathy was induced by performing a Chronic Constriction Injury (CCI) of the sciatic nerve, as previously described^24^. Briefly, under isoflurane (2%) anesthesia the skin on the lateral surface of the thigh was incised and a section made directly through the biceps femoris muscle exposing the sciatic nerve. Three ligatures (10-0 prolene) around the nerve were used until they elicited a brief twitch in the hindlimb. Subsequently, the skin was sutured with a non-absorbable 6-0 polypropylene suture, and local asepsis was performed using a 10% povidone-iodine® solution. Sham-operated were used as a control.

#### Nociceptive behaviors tests

All behavioral tests on mice were performed after a minimum of 30 minutes of habituation to the environment and assays.

#### von Frey filament test

For testing mechanical nociceptive threshold, mice were placed on an elevated wire grid and the plantar surface of the ipsilateral hind paw stimulated perpendicularly with a series of von

Frey filaments increasing logarithmically in force applied. Mechanical pain hypersensitivity (mechanical allodynia) was examined by measuring paw withdrawal thresholds in the lateral area of the hind paw, corresponding to the sural nerve innervation – the branch of the sciatic nerve preserved in the SNI model. Each von Frey hair filament was applied for a maximum of 3 seconds and paw withdrawal recorded. The data collected was done considering the ascending method - the response was with a series of von Frey filaments, with logarithmically increasing stiffness (0.008–2.0 g)^69^.

For the frequency of response analyses, the number of animal responses to 10 consecutive paw stimulations with 0.02 or 0.008 filaments depending on the experiment were determined^5,70^ Clique ou toque aqui para inserir o texto..

#### Electronic von Frey test

Using the same elevated wire grid described, a hind paw flexion reflex was evoked with a hand-held force transducer, adapted with a 0.5-mm^2^ polypropylene tip. The stimulus was applied in the hind paw, with a gradual increase in pressure. When the paw was withdrawn, the stimulus was automatically discontinued, and its intensity recorded. The end point was characterized by the removal of the paw in a clear flinch response after the paw withdrawal^69^ .

#### Hargreaves test

The latency of paw withdrawal to radiant heat stimuli was measured using the Hargreaves apparatus and described previously^71^. Mice were habituated in an elevated glass surface of constant temperature (32°C). An infrared light source was applied perpendicularly to the hind paw plantar surface of each mouse. The end point was characterized by the removal of the paw followed by clear flinching movements. Latency (time in seconds) to paw withdrawal was automatically recorded. To avoid tissue damage, a maximum latency (cut-off) was set at 20 seconds.

#### Hot-plate test

Noxious heat thresholds were also examined using the Hot-Plate test. Mice were placed in a 10-cm-wide glass cylinder on a hot plate maintained at 48°C, 52°C or 56°C. Mice were each tested twice at least 20 minutes apart. The latencies in seconds for hind paws licking or jumping for each animal were recorded. To avoid tissue damage, a maximum latency (cut-off) was set at 20 seconds^70^.

#### Acetone test

Mice were habituated in the same cages and elevated grids used during the von Frey tests. Then, a drop (50 µl) of acetone was applied to the plantar surface of the hind paw using a syringe of 1 ml. The time spent with nociceptive behaviors - defined as flinching, licking or biting the limb - were timed within 60 seconds after the application of acetone^70^.

#### Rotarod test

The apparatus consisted of a bar with a diameter of 2.5 cm, sub-divided into five compartments by disks 25 cm in diameter. The bar rotated at a constant speed of 22 rotations per minute. All animals were trained on the rotarod, with only those animals able to remain on the bar for two consecutive periods of 120 seconds selected for subsequent testing 24 hours later. Mice were submitted to walk on the rotating cylinder for a total of 120 seconds. The time that each animal remained walking was registered as latency to fall^70^.

#### Intrathecal and oral treatments

Under isoflurane (2%) anesthesia, mice were securely held in one hand at the pelvic girdle, and a BD Ultra-Fine® (29G) insulin syringe was inserted directly into the subarachnoid space (close to L4–L5 segments) of the spinal cord. A lateral movement of the tail indicated proper placement of the needle in the subarachnoid space^72^. For all deliveries, we used 5-7 µl of volume. In all experiments in which intrathecal injections were performed, the vehicle was replaced by aCSF (artificial cerebrospinal fluid; Tocris, Catalog number: 3525). The i.t route was used to deliver FITC-albumin (5 μg/site/5 μL, Sigma-Aldrich) into the subarachnoid space (SAS) to track molecules diffusing into the cerebrospinal fluid (CSF). To study exchange of cells between bone marrow and SAS, the CXCR4 inhibitor, on days 7 and 9, two groups of mice (control or SNI) received vehicle and two groups (control or SNI) were treated intrathecal with AMD3100 (5 µg/site). On days 6, 7, 8, and 9, two groups of mice (control or SNI) received vehicle and two groups (control or SNI) received intrathecal CHZ868 (20 µg/site), a type II JAK2 inhibitor, every 24 h. During the 3-day rmGM-CSF treatment period, two groups of mice (control or SNI) received vehicle, whereas two other groups (control or rmGM-CSF) were orally treated with Fedratinib (60 mg kg^-1^, twice daily, a type II JAK2 inhibitor,) by gavage, administered 1 h before rmGM-CSF injection, with a 12 h interval between doses. To study the pro-nociceptive effect of meningeal GM-CSF and mechanisms underlying, rmGM-CSF (100 ng/site/5 μL) was injected intrathecal twice a day, 12 h apart followed by the pain behavior analysis, quantification of meningeal leukocytes and vertebral BM emergency myelopoiesis.

#### Chemicals and reagents

AMD3100 octahydrochloride (Abcam, Catalog number: ab120718), CHZ868 (MedChemExpress, Catalog number: HY-18960) and Fedratinib (MedChemExpress, Catalog number: HY-10409). For mouse *in vivo* studies and in vitro electrophysiology studies using mouse DRG neurons, recombinant mouse GM-CSF (R&D Systems, Catalog number: 415- ML) was used at the indicated concentrations and doses. For in vitro electrophysiology studies using rat DRG neurons, recombinant rat GM-CSF (R&D Systems Catalog number:518-GM) was used. In addition, inflammatory mediators ATP (Catalog number: A2383), bradykinin (Catalog number: B3259), histamine (Catalog number: H7125) and noradrenaline (Catalog number: A7257) were all purchased from Sigma-Aldrich; prostaglandin E_2_ was purchased from Acros Chemicals (Catalog number: 233560100). Drugs were prepared immediately before use and antibodies were kept in ice until injection. Systemic drugs were prepared in sterile PBS pH 7.4 (Gibco, Catalog number:10010023), while intrathecal drugs were diluted in aCSF (artificial cerebrospinal fluid) (Tocris, Catalog number: 3525).

#### Dorsal root or brain meninges collection for Real-Time RT-PCR and Flow Cytometry

Mice were terminally anesthetized with xylazine (10 mg kg^−1^) and ketamine (100 mg kg^−1^) by intraperitoneal (i.p.) injection followed by intracardiac perfusion with phosphate-buffered solution (0.01 M PBS) (20 ml min^-1^ for 2 minutes) at room temperature to wash the blood and consequently the circulating leukocytes. For dorsal root meninges, we realized a laminectomy to remove vertebral arcs in lumbar segments between L3 and L5. Vertebral arcs were removed and DRG and spinal cord were exposed carefully to preserve the external surface of meninges covering both tissues. Then, DRG’s meninges (from L3, L4 and L5 DRG) on the ipsilateral side were removed (peel off) and pulled towards the spinal cord meninges. The DRG were then discarded. Next, the meninges were cut vertically just below the spinal cord on the ipsilateral side and vertically in the spinal cord on the contralateral side, just before reaching the DRGs. Finally, the meninges were pulled in a caudal-to-rostral (L5-L3) direction.

To collect the brain meninges, first, the skullcap was removed, then, the brain meninges were peeled and carefully removed from the inside of the skullcap with fine forceps as previously described^30^.

#### Real-time RT-PCR of dorsal root meningeal tissue or vertebrae BM

Dorsal root meninges were collected from mice as described in the “*Dorsal root and brain meninges collection for Real-Time RT-PCR and Flow Cytometry.* Regarding the vertebrae BM, the ipsilateral side of L3, L4 and L5 were collected. The samples, once collected, in a Trizol reagent (Invitrogen, Catalog number: 15596018), were immediately stored on dry ice and later in a freezer at -70°C. After grinding the tissue, the extraction of the total RNA was carried out according to the reagent manufacturer, namely: separation in chloroform, precipitation in isopropanol (overnight -20°C) and washing with pure ethanol. The RNA concentration of each sample was determined by the optical density at the wavelength of 260 nm (NanoDrop, ThermoFisher, USA). One μg or 500 ng of total RNA was transcribed to complementary DNA by the action of the MultiScribe® reverse transcriptase enzyme, high- capacity kit (Life-Invitrogen, Cat. 4368814). Sequence of primers used are: *Csf2* - Forward: CCTGGGCATTGTGGTCTACAG / Reverse: GGCATGTCATCCAGGAGGTT; *Cd19* - Forward: TCATTGCAAGGTCAGCAGTGT; Reverse: GGAGAGCACATTCCCGTACT; *Cxcl12* - Forward: ATCAGTGACGGTAAACCAGT; Reverse: TCTGTTGTTGTTCTTCAGCC. RT-PCR was performed with a final reaction volume of 10 µl, SYBR-Green. The reaction was performed using the QuantStudio5 Real-Time PCR System (Life-Applied Biosystems). Results were analyzed using the comparative method of "cycle threshold" (CT).

### Histology

#### Whole-mount meningeal and whole-mount DRG tissue for immunofluorescence and confocal imaging

At 10 post-SNI, CCI or sham surgeries, mice were terminally anesthetized with xylazine (10 mg kg^−1^) and ketamine (100 mg kg^−1^) by intraperitoneal (i.p.) injection followed by intracardiac perfusion with phosphate-buffered solution ice-cold (1X PBS) (20 ml min^-1^ for 2 minutes) and 4% paraformaldehyde (PFA) (20 ml min^-1^ for 4 minutes) (Sigma Aldrich, Catalog number: n. 158127). After fixation, we removed the dorsal muscles to gain access to the vertebrae. Then we separated the gliding joints between the vertebral zygapophyses, and a laminectomy was performed to remove the vertebral arches along the lumbar segment. The laminectomy was carefully performed to preserve the tissue architecture of the meninges covering the dorsal surface of the spinal cord and DRGs.

The entire meninges covering the lumbar spinal cord segment were carefully collected, preserving their dorsal and lateral portions. Of note, samples were collected without removing the attached sensory ganglia along the lumbar segment. Using a stereomicroscope (LEICA, EZ4 W), the remaining spinal cord and tracts were carefully removed without disrupting the meningeal tissue and post-fixed in 4% PFA for 2 to 24 hours. The whole-mount samples were then washed 3 times for 10 minutes in PBS with 0.4% Triton X-100 for 10 minutes, permeabilized with 0.1% Triton X-100 for 24 hours (Millipore Sigma, Catalog number: T9284) and blocked and permeabilized in PBS with 0.4% Triton X-100, 0.2M of glycine and 2% bovine serum albumin or 5% goat serum for 4 hours (Invitrogen, Catalog number: 50062Z) for 60 minutes at room temperature. Next, the samples were incubated with primary antibodies for 24 hours at 4°C. Then, the samples were, washed 3 times for 5 minutes each time in PBS with 0.4% Triton X-100 at room temperature, and incubated with secondary antibodies together with DAPI (Catalog Number: D3571) in PBS with 0.4% Triton X-100, 0.2M of glycine and 2% bovine for 120 minutes at room temperature. Finally, the samples were washed 3 times for 5 minutes each time in PBS with 0.4% Triton X-100 and mounted for confocal microscopy. Whole mount meninges and associated ganglia were carefully positioned on glass coverslip and coverslipped with Fluoromount-G™ Mounting Medium (Invitrogen, Catalog Number: 00-4958-02) or FluorSave (Millipore Sigma, Catalog number: 345789) and a glass coverslip. The same procedure was used for the L4 lumbar DRG whole mounts. The following antibodies were used: anti-CD45 (1:500; Biolegend, Clone 30F-11, Catalog Number: 103102), anti-CD31 (1:200; Millipore Sigma, Clone 2H8, Catalog Number: MAB1398Z), Full-Length anti-ZsGreen (1:500; Takara, Polyclonal, Catalog Number: 632474), anti-F4/80 (1:250; Cell signaling Technology, Catalog Number: 71299), anti-Ly6G (1:300; Abcam, Catalog Number: AB25377), anti-Podoplanin (1:350; Invitrogen, Catalog Number: 14-5381-82). The dilution of antibodies was always twice the primary antibody used.: Alexa fluor 594 (Catalog number: A11007), Alexa fluor 488 (Catalog number: A11006), Alexa fluor 488 (Catalog number: A21110) or Alexa fluor 647 (Invitrogen, Catalog number: A21472).

Meningeal whole mounts and DRG whole mounts were imaged using either a Zeiss LSM780 or Nikon Eclipse Ti2 microscope using 10× or 40× objectives (pinhole 1 AU, 1024×1024 pixels, 1.5 µm z-steps). After acquisition, images were processed using the FIJI package for ImageJ software.

### Light-sheet microscopy imaging and analysis

#### Tissue preservation and clearing, immunolabeling and imaging

Clearing, immunolabeling and imaging of spinal cord vertebral segments were performed by LifeCanvas Technologies through a contracted service. Paraformaldehyde-fixed tissue samples were preserved using SHIELD reagents^73^ according to the manufacturer’s protocol (LifeCanvas Technologies). Preserved samples were subsequently decalcified in 10% EDTA (in deionized water) at 4°C for 7 days. Delipidation was performed in two steps. First, organic delipidation was carried out using a modified FDISCO protocol^74^, with samples incubated for 1 hour in each serial wash steps at 4°C and 2 days incubation in 100% dichloromethane (DCM) at 4°C. This was followed by aqueous delipidation using Clear+ reagents (LifeCanvas Technologies) for 7 days at 45°C.

Immunolabeling was conducted using the SmartBatch^+^ system and RADIANT buffer (LifeCanvas Technologies), incorporating stochastic electrotransport^75^ and eFLASH/CuRVE technologies^76^. The primary antibody used was anti-CD45 (1:450; R&D Systems, Catalog Number: AF114). The secondary antibodies SeTau-647 (1:1500; Canvas Technologies, Catalog Number: DkxGt-ST). Following labeling, samples were incubated in 50% EasyIndex (diluted in deionized water) overnight at 37°C, then transferred to 100% EasyIndex (RI = 1.52; LifeCanvas Technologies) for 24 hours for refractive index (RI) matching. After RI matching, samples were imaged using a SmartSPIM axially swept light-sheet microscope equipped with a 3.6× objective (0.2 NA; LifeCanvas Technologies). Images were analyzed with Imaris software (10.2.0) using the 3D View and animations were generated by adding an ortho slicer solid object. For videos, a sequence of frames was captured while rotating the image object along the anterior–posterior axis of the spine.

#### FDISCO processing

All procedures were carried out at 4 °C with shaking as previously described^74^. Fixed samples were washed in 50%, 70%, and 80% tetrahydrofuran (THF) in 25% quadrol (in 1× PBS to adjust to pH 9), then 95% THF twice. Timing of organic delipidation washes followed the outline given below, with 3-week-old juvenile whole-body samples treated with dichloromethane 3 times. Samples were delipidated with 100% dichloromethane (DCM; Sigma-Aldrich) then washed in 95% THF twice, followed by 80%, 70%, and 50% THF. Samples were then washed in 0.2% PBST for 1hour then 2hours at RT, followed by 1× PBS for 1hour 3 times then overnight at RT to thoroughly wash out any remaining organic solvent.

#### Immunofluorescence of DRGs and spinal cord

Mice were terminally anesthetized with xylazine (10 mg kg^−1^) and ketamine (100 mg kg^−1^) by intraperitoneal (i.p.) injection followed by intracardiac perfusion with phosphate-buffered solution (0.01 M PBS) (20 ml min^-1^ for 2 minutes) at room temperature. The samples, once collected, were immediately stored in cold 4% PFA and kept at 4°C for 30 minutes (DRG) and 2 hours for (spinal cord) and then transferred to the 30% sucrose solution for 2 days at 4°C. On dry ice, tissues were embedded in TissueTek® OCT Compound (Sakura, Catalog number: 4583) blocks and cut on the cryostat: 14 μm for DRG and 40 μm for spinal cord. In well plates or slides, the samples were washed with a PBS buffer, incubated in blocking solution (2% BSA, 22.52 mg ml^-1^ glycine) in 0.01M PBS, both at room temperature for 1 hour. Posteriorly, tissues were incubated overnight with the primary antibody at 4°C and after, for 2 hours with the secondary antibody coupled to fluorophore. The following antibodies were used: anti-Iba1 (1:300; Wako, Polyclonal, Catalog Number: 016-20001), Full-Length anti-ZsGreen (1:500; Takara, Polyclonal, Catalog Number: 632474), β-III-Tubulin/Tuj1 (1:25; R&D Systems, 637-conjugated monoclonal, Catalog Number: NL1195V). The dilution of antibodies was always twice the primary antibody used.: Alexa fluor 594 (Catalog number: A11007), Alexa fluor 488 (Catalog number: A11006), or Alexa fluor 647 (Invitrogen, Catalog number: A21472). The slides were assembled using VectaShield® with DAPI (Vector, Catalog number: H-1200). Images were acquired using a confocal Zeiss fluorescence microscope (AxioObserver LSM780) and images were analyzed using NIH Image J (Fiji) open-source software.

#### Spine column sections for immunofluorescence and confocal imaging

Mice were terminally anesthetized with xylazine (10 mg kg^−1^) and ketamine (100 mg kg^−1^) by intraperitoneal (i.p.) injection followed by intracardiac perfusion with phosphate-buffered solution (0.01M PBS) (20 ml min^-1^ for 2 minutes) at room temperature. This procedure was followed by additional perfusion with 4% paraformaldehyde (4% PFA) (Sigma Aldrich, Catalog number: 158127) for tissue fixation (20 ml min^-1^ for 3 minutes). Next, the lumbar segment of the spinal column was removed from neuropathic or control *Ccr2*^RFP^/*Cx3cr1*^eGFP^ mice, post-fixed in 4% PFA for 24 hours, and transferred to decalcification solution (PBS in 10% EDTA). Samples were left in the decalcification solution, which was renewed daily, with gentle and constant agitation at room temperature (24°C) for 14 days. The samples were immersed in PBS with sucrose (10% and 20 % for 12 hours and 30% overnight, sequentially) and then embedded in a medium for frozen tissue (Tissue-Tek O.C.T. compound) and frozen over dry ice. Samples were sliced in the transverse plane (sections with 30μm) on a cryostat (Leica). Sections were then washed in PBS with 0.4% Triton X-100 for 10 minutes, blocked and permeabilized in PBS with 0.2% Triton X-100 and 2% bovine serum albumin for 60 minutes at room temperature. Next, the samples were incubated with primary antibodies for 24 hours at 4°C, washed 3 times for 5 minutes in PBS with 0.2% Triton X-100 at room temperature, and incubated with secondary antibodies in PBS with 0.4% Triton X-100 and 2% bovine for 120 minutes at room temperature. Immunostained samples mounted on Starfrost slides were coverslipped with Vectashield Antifade Mounting Medium with DAPI (Vector Laboratories, Catalog number: H-1200) and a glass coverslip. Sections were imaged using a Zeiss LSM780. After acquisition, images were processed using the FIJI package for ImageJ software.

#### Laminectomized spine column samples for scanning electron microscopy (SEM)

Mice were terminally anesthetized with xylazine (10 mg kg^−1^) and ketamine (100 mg kg^−1^) by intraperitoneal (i.p.) injection followed by intracardiac perfusion with phosphate-buffered solution (0.0.1M PBS) at room temperature (20 ml min^-1^ for 2 minutes). This procedure was followed by additional perfusion with 4% paraformaldehyde (4% PFA) (Sigma Aldrich, Catalog number: 158127) for tissue fixation (20 ml min^-1^ for 3 minutes). After fixation, we removed the dorsal muscles to gain access to the vertebrae. Then we separated the gliding joints between the vertebral zygapophyses, and a laminectomy was performed to remove the L4 vertebral arch. The laminectomy was carefully performed to preserve the tissue architecture of the meninges covering the dorsal surface of the spinal cord and sensory ganglia. From this boneless sample, the L4 segment containing the right and left dorsal root ganglia (DRG) was separated and prepared for scanning electron microscopy (SEM).

Two steps were used to fix the samples. First, the laminectomized spine was immersed in a 2.5% glutaraldehyde buffer for fixation. Next, the samples were immersed in 1% osmium tetroxide solution and subsequently dehydrated through an ethyl alcohol gradient (30%, 50%, 70%, 90% and 95% for 10 minutes and 3 times at 100% for 20 minutes). The spinal dehydrated pieces were then dried with CO2 using Critical Point Dryers (CPD 030 Bal-Tec) and metalized with gold using a sputtering device for SEM coating (SCD 050 Sputter counter Bal-Tec). Finally, the samples were properly mounted in a SEM specimen stub with an appropriate coal glue and forwarded for scanning. It is noteworthy that the specimens were carefully positioned in the stubs so that the laminectomized dorsal aspect of the specimens was exposed for subsequent scanning. The samples adhered on the top of the SEM specimen stub were imaged using the MEV JSM-6610LV (JEOL).

#### Microtomography (μCT) of the mouse or human lumbar vertebrae samples

C57BL/6J mice were euthanized and the entire lumbar segment without skin was removed. The samples were then fixed for 1 week and scanned using the SkyScan 1172 system (Bruker, Kontich, Belgium) with a 100 kV X-ray source detected by an 11-megapixel camera with a resolution of up to 1 μm. The NRecon Cluster software (version 1.15.4.0, Micro Photonics Inc., Allentown, PA, USA) was used to obtain two-dimensional tomographic projections and three-dimensional (3D) reconstructions.

To track vertebral ossified channels in humans, two lumbar spines from human cadavers previously preserved in formaldehyde were used. These samples were sourced ethically, and their research use was in accord with the terms of the informed consents under an IRB/REC approved protocol. Sectioned vertebral archs from lumbar vertebra were then prepared for scanning. Samples were imaged using a SkyScan 1172 microCT system (Bruker, Kontich, Belgium) equipped with an 11-megapixel Hamamatsu C9300 camera. Scans were acquired with an isotropic voxel size of 9.9 µm, at 89 kV tube voltage, 112 µA tube current, 1240 ms integration time, and 360 projections. Image reconstruction was performed using NRecon software (v1.6.10.4, Bruker) with smoothing set to 2, ring artifact correction to 6, and beam hardening correction to 20%. Linear measurements of bone channels were obtained along all three spatial axes (3D). The maximum diameter of each segmented channel was determined manually using DataViewer software (v1.5.6.2, 64-bit). Calibration was based on the attenuation coefficient.

### Flow cytometry analyses

#### DRG’s and brain meningeal cells

*DRG’s* or brain meninges were collected from mice as described in the “*DRG’s meninges collection for Real-Time RT-PCR and Flow Cytometry***”**. Samples were immediately placed in 500 µl of cold DMEM High Glucose (Corning, Catalog number: 10013-CV) supplemented with 10% Fetal Bovine Serum (FBS) (Gibco, Catalog number: 12657-029) in a 2 ml Eppendorf tube. The tissue samples were finely minced with scissors inside the Eppendorf tube. Then, 500 µl of a solution containing 1 mg ml^-1^ of Collagenase Type D (Sigma Aldrich, Catalog number: 11088866001 and 0.1 mg ml^-1^ of DNase Type I (Sigma Aldrich, Catalog number: DN25) was added, and the sample was incubated at 37°C under constant agitation at 600 rpm in a thermoblock for 45 minutes. After incubation, the samples were homogenized and immediately added 1 ml of DMEM High Glucose (Gibco, Catalog number: 12657-029) supplemented with 20% FBS (Gibco, Catalog number: 12657-029). The digested tissue was passed through a 40-70 µm cell strainer (Corning, Catalog number: 431750; Miltenyi Biotec, Catalog number: 130-095-823 using a 1 ml syringe plunger directly into the FACS tube (Falcon, Catalog number: 352008) and then centrifuged at 450 g for 5 minutes at 4°C. Blood was collected by retro-orbital bleeding under anesthesia at the time the mice were killed. Samples were centrifuged for 10 minutes at 450 g, and red blood cell lysis was performed with ammonium-chloride-potassium (ACK) lysis buffer (Quality Biological) for 7 minutes. Samples (meninges or blood) were washed in the Running buffer (Milteney Biotec, Catalog number: 130-091-221) and the supernatant was carefully discarded using a pipette. The cells obtained were resuspended in 30 µl of Running Buffer (MACS, Catalog number: 130-091- 221) containing specific monoclonal antibodies against surface markers for 30 min on ice. Dead cells were excluded by Fixable Viability Dye (1:3000; eBioscience, Catalog Number 65-0865-14). The following monoclonal antibodies were used: anti-CD45-PE Cy 7 (1:200; Biolegend, Clone 30F-11, Catalog Number: 103113), anti-CD11b-APC (1:200; BD Biosciences, Clone M1/70, Catalog Number: 561690C), anti-CD11b-FITC (1:200; eBioscience, Clone M1/70, Catalog Number: 11-0112-82), anti-Ly6G-BV510 (1:200; BioLegend, Clone 1A8, Catalog Number: 127633), anti-Ly6G-PE (1:200; BD Biosciences, Clone 1A8,Catalog Number: 551461), anti-Ly6C-PERCP Cy5.5 (1:300; eBioscience, Clone HK1.4, Catalog Number: 45-5932-82), anti-CD11c-PE (1:100; Biolegend, Clone N418, Catalog Number: 117308), anti-CD11c-BV421 (1:100; Biolegend, Clone N418, Catalog Number: 117329), anti-CD3-FITC (1:200; eBioscience, Clone 145-2C11, Catalog Number: 11-0031-82), anti-CD4-BUV605 (1:200; BioLegend, Clone RM4-5, Catalog Number: 100548) and anti-CD8-BV421 (1:200; BioLegend, Clone 53-6.7, Catalog Number: 100737). Then, 500 microliters of Running Buffer were pipetted and centrifuged for 5 minutes at 450 g. The supernatant was removed, and each eppendorf tube was resuspended with 200 microliters of Running Buffer and added 20% AccuCheck Counting Beads (Thermo Fisher, Catalog number: PCB100). Samples acquisition was performed by the FACSCanto™ II flow cytometry instrument (BD Biosciences, San Jose, CA, USA) or FACSymphony™ A1 flow cytometry instrument (BD Biosciences, San Jose, USA) and data were analyzed using FlowJo software (Treestar, Ashland, OR, USA).

#### Vertebra or tibia BM cells

The vertebrae (L3, L4 and L5) or tibia ipsilateral to the SNI surgery were collected immediately after perfusing the mouse with ice-cold 0.01M PBS, pH 7.2. The tissue was immediately placed in 500 µl of cold DMEM High Glucose (Corning, Catalog number: 10013-CV) supplemented with 10% Fetal Bovine Serum (FBS) (Gibco, Catalog number: 12657-029) in a 2 ml Eppendorf tube. Then, they were triturated using scissors and subsequently transferred with a Pasteur pipette to a mortar, where they were mechanically triturated with a pestle. Next, the sample was passed through a 70 µm (Corning, Catalog number: 431751) sieve into a Petri dish using a plunger. The entire volume was then passed through a 100 µm (Corning, Catalog number: 431752) sieve into a 2 ml Eppendorf tube and centrifuged for 10 minutes at 450 g. The cells obtained were resuspended in 100 µl of Running Buffer containing specific monoclonal antibodies against surface markers for 30 min on ice. Dead cells were excluded by Fixable Viability Dye (Catalog Number 65-0865-14, eBioscience, 1:3000). The following monoclonal antibodies were used: anti-CD45-PE Cy7 (1:200; Biolegend, Clone 30F-11, Catalog Number: 103113), anti-CD11b-APC (1:200; BD Biosciences, Clone M1/70, Catalog Number: 561690C), anti-CD11b-FITC (1:200; eBioscience, Clone M1/70, Catalog Number: 11-0112-82), anti-Ly6G-BV510 (1:200; BioLegend, Clone 1A8, Catalog Number: 127633), anti-Ly6G-PE (1:200; BD Biosciences, Clone 1A8, Catalog Number: 551461), anti-Ly6C-PERCP Cy5.5 (1:300; eBioscience, Clone HK1.4, Catalog Number: 45-5932-82), anti-CD11c-PE (1:100; Biolegend, Clone N418, Catalog Number: 117308) and anti-CD11c-BV421 (1:100; Biolegend, Clone N418, Catalog Number: 117329). Cells were washed and resuspended with 200 µl Running buffer (Miltenye Biotec, Catalog number: 130-091-221) adding 20% AccuCheck Counting Beads and transferred to a FACS tube with filter (Falcon, Catalog number: 352225). For CMP and GMP analysis, cells were stained with antibodies against: ckit-BV510 (BioLegend, clone 2B8, 1:100; Catalog Number: 105839), Sca1-PECy7 (BioLegend, clone D7, 1:100; Catalog Number: 108114) and Lineage (Lin)- Pac blue (BioLegend, 1:200; Catalog,Number: 133310), CD34 (BD Biosciences, clone, BD, RAM34, 1:100; Catalog Number: 551387) and CD16/32 (BioLegend, clone 2.4G2, 1:100; Catalog Number: 561728). Antibodies listed were incubated for 30 minutes on ice. Cells were further stained with the Click-iT Plus EdU Cell Proliferation Kit (Invitrogen™, Catalog number: C10640) following the manufacturer’s guidelines for analysis. For CMP and GMP proliferation, mice received injection of EdU (25 mg kg^−1^, i.p.)^30^. 24h before samples collection. For rmGM-CSF, mice received three injections of EdU (25 mg kg^−1^, i.p.) 1 hour after each rmGM-CSF injection, 24 hours apart and were sacrificed 24 hours after the final injection. Cells were washed and resuspended with 200 µl Running buffer adding 20% AccuCheck Counting Beads (Thermo Fisher, Catalog number: PCB100). Sample acquisition was performed by the FACSCanto™ II flow cytometry instrument (BD Biosciences, San Jose, CA, USA) or FACSymphony™ A1 flow cytometry instrument (BD Biosciences, San Jose, CA, USA) and data were analyzed using FlowJo software (Treestar, Ashland, OR, USA).

#### Fluorescence-activated cell sorting (FACS) of DRG’s meningeal leukocytes

Samples from the dorsal root meninges were dissected and processed as previously described in “*Flow Cytometry analyses of DRG’s and brain meningeal cells*” Primary mouse CD45^+^ cells were purified from the dorsal root meninges of mice using a FACSAriaIII sorter (BD Biosciences, Milpitas, CA, USA). Cell purity was confirmed to be ≥ 95%.

#### scRNA-seq of leukocytes from DRG’s meninges

Cell suspension samples containing *DRG’s* meningeal immune cells (CD45^+^) were obtained as described in the section “*Fluorescence-activated cell sorting (FACS) of DRG’s meningeal cells*” and used for droplet-based scRNA-seq (10x Genomics). The quality of the single-cell suspensions (viability >90%) was verified immediately before encapsulation performed in a Chromium Controller instrument (10X Genomics, Catalog Number: PN120270). The single cell libraries were prepared using the Chromium Next GEM Single Cell 3′ Reagent kit v3.1 (10x Genomics, Catalog Number: PN1000269) and the Gene Expression v3.1 dual index protocol (10x Genomics, Document Number: CG000315 Rev E). Following encapsulation and mRNA barcoding, cDNA was synthesized, isolated and amplified following the manufacturer’s instructions. The quality of the amplified cDNA (concentration, size and purity) was verified using a High Sensitivity D1000 ScreenTape Assay, Catalog Number: PN- 5067-1504) and 4200 TapeStation system (Agilent Technologies, Catalog Number: G2991). Next, amplified cDNA was submitted to fragmentation, purification (DNA size selection and cleanup), end repair, A-tailing, adaptor ligation and sample index PCR amplification following the manufacturer’s instructions. After library construction, quality control (concentration, size and purity) was performed using a High Sensitivity D1000 ScreenTape Assay, Catalog Number: PN- 5067-1504) and 4200 TapeStation system (Agilent Technologies, Catalog Number: G2991). For sequencing was used the HiSeq 4000 Kit (150 cycles) (Illumina, Catalog Number: PE-410-1001, FC-410-1002) and. The CD45^+^ sample libraries were pooled and equally distributed in flow cell (Read 1: 28 bp; i7 index: 10 bp; i5 index: 10 bp; Read 2: 90 bp) and runs on the Illumina Hiseq 4000 System (Illumina, Catalog Number: SY-401-4001). Library preparation and sequencing were conducted by the Hemocenter Sequencing Core facility at Ribeirao Preto Medical School.

CellRanger filtered cells and counts were used for downstream analysis in Seurat version 5.3.0 implemented in R version 4.4.1. Cells were excluded if they had fewer than 200 features, more than 7500, or the mitochondrial content was more than 15%. For control and post-injury samples, reads from multiple samples were integrated and normalized following a standard Seurat CCA integration pipeline^77^. PCA was performed on the top 3000 variable genes and the top 30 principal components were used for downstream analysis. A K-nearest- neighbor graph was produced using Euclidean distances. The Louvain algorithm was used with a resolution set to 1.2 to group cells together. Nonlinear dimensionality reduction was done using UMAP. Zero imputation was performed using the Alra pipeline in the SeuratWrappers package^78^. Cell types were defined by comparing cluster marker genes with previously published scRNA-seq datasets of mouse meninges^79,80^. Visualization of genes illustrating expression levels was performed using R/Seurat commands (DimPlot, FeaturePlot, and DotPlot) using ggplot2^81^, Nebulosa^82^ R packages.

#### Pathway enrichment analysis

To evaluate the enriched pathways in the clusters, we used the Seurat FindAllMarkers function and the Wilcoxon rank-sum test, to obtain the differentially expressed genes in both control and post-SNI injury samples and submitted these genes to clusterProfiler package^83^. Gene ontology (GO) terms in the Biological Processes category with *P* < 0.05 were considered significant. Statistically significant, non-redundant GO-enriched terms were plotted.

#### Ligand-receptor interaction analysis with CellChat

Cell-cell communications were analyzed by applying the R package CellChat developed previously^84^. Data obtained from scRNA-seq was processed and the CellChat object was initialized. Next, the ligand-receptor interaction database was set and the expression data was further preprocessed for cell-cell communication analysis. Afterward, the communication probability was computed, and the cellular communication network was inferred.

#### Spatial transcriptome from human DRGs with painful diabetic neuropathy (PDN)

All human tissue collection and experimental procedures were conducted in compliance with institutional ethical guidelines and were approved by the Institutional Review Board (IRB) of The University of Texas at Dallas (Protocol: Legafavy MR-15-237). Lumbar dorsal root ganglia (DRGs) were obtained from organ donors through the Southwest Transplant Alliance (STA), an accredited organ procurement organization (OPO) in Texas. A secure informed consent for research use of donor tissues either directly from the individual donor or from the donor’s next of kin. DRGs were dissected through a ventral surgical approach as described previously^38,85^. Spatial transcriptomic profiling was performed using the 10x Genomics Visium v1 platform following the manufacturer’s instructions and previously published protocols^86^. Tissue sections were visualized and analyzed in Loupe Browser (10x Genomics). Spots exhibiting overlapping expression of canonical ILC2 markers (KLRB1, IL1RL1, PTPRC, GATA3) in the absence of pan–T cell markers (CD3E, CD3D, CD4, CD8A) were identified for each sample. Quantification of ILC2-associated barcodes was performed, results were plotted, and statistical analyses were run using GraphPad Prism (version 10.6.1).

#### Generation of bone marrow-derived macrophages (BMDMs)

BMDMs were obtained using L929 cell-conditioned medium, as previously described^87,88^. Bone marrow cells were collected from femurs and differentiated with RPMI 1640 medium (Corning, Catalog number: 15-040-CV), supplemented with 20% L929 cell-conditioned media, 20% FBS, L-glutamine (2 mM, Globo, Catalog number: 35050061), penicillin (100 U ml^-1^ 2.5 µg ml^-1^, Sigma Aldrich, Catalog number: P4333) and fungizone (2.5 µg ml^-1^, Gibco, Catalog number: 15290-018) for 7 days at 37°C and 5% CO2. Differentiated BMDM (>95% of cells CD11b^+^ F4/80^+^) were collected and seeded in 96-well culture plates at density of 2 x 10^5^ cells/well and kept in RPMI 1640 complete media, 10% FBS, L-glutamine (2 mM), penicillin (100 U ml^-1^) and fungizone (2.5 µg ml^-1^) for 2 h. BMDMs were treated with recombinant mouse GM-CSF (20 ng ml^-1^), IL-4 (20 ng ml^-1^) or LPS (100 ng ml^-1^) for 24 h at 37 °C and 5% CO_2_. Culture supernatant was collected for cytokine and chemokine quantification by ELISA.

#### Enzyme-linked immunosorbent assay (ELISA)

Concentrations of CCL17, CCL22, CCL2 and TNF in BMDM culture supernatants were determined by ELISA according to the manufacturer’s instructions (R&D Systems). Absorbance was measured at 450 nm on the SpectraMax 190 Microplate Reader spectrophotometer using the SoftMax Pro software.

### Electrophysiology

#### Isolation and primary culture of mouse DRG

Mice were sacrificed by rising concentrations of CO_2_ followed by exsanguination. The spinal column was removed, split bilaterally, and the DRG dissected free before placing in ice-cold Leibovitz L-15 Glutamax media. Roots were trimmed and DRG washed in ice-cold Dulbecco’s Phosphate Buffered Saline (without MgCl_2_ and CaCl_2_; DPBS) prior to enzymatic dissociation in L-15 media supplemented with 24 nM NaHCO_3_ and 6 mg/ml bovine serum albumin solution, firstly with collagenase (15 mins at 37 °C & 5 % CO_2_; 1 mg ml^-1^; Type 1A, Sigma-Aldrich C9891) then secondly with trypsin (30 mins at 37°C & 5% CO_2_; 1mg ml^-1^; Sigma-Aldrich T9935). DRG were subsequently mechanically dissociated by trituration with a pipette before resuspension in growth media (Lebovitz L-15 Glutamax media supplemented with 10 % FCS, 24 mM NaHCO_3_, 38 mM glucose and penicillin/streptomycin). Cells were plated in growth media onto 12mm glass coverslips precoated with PDL and laminin and incubated at 37°C and 5 % CO_2_ until use.

#### DRG neuron patch-clamp electrophysiology

Whole-cell recordings of primary cultured mouse DRG neurons were made at room temperature (∼22C) using a MultiClamp 700B amplifier (Molecular Devices) connected to a DigiData 1550A interface (Molecular Devices) controlled by ClampEx11.1 software (Molecular Devices) with a 10kHz acquisition rate. Patch electrodes were pulled (Narishige PC-100), heat polished (Narishige MF-83) and filled with intracellular solution for current-clamp recordings (ICS; in mM: KCl 140, EGTA 0.5, HEPES 5, Na2ATP 3; adjusted to pH7.3 with KOH and ∼300 mOsm with glucose) yielding typical pipette resistances of 2-3 MΩ. On formation of whole-cell configuration access resistances were generally < 8 MΩ. Cells were continuously superfused with extracellular solution (ECS*_patch_*; in mM: NaCl 140, KCl 3, MgCl_2_ 2, CaCl_2_ 2, HEPES 10; adjusted to pH7.3 with NaOH and ∼315 mOsm with glucose) by gravity-fed rapid exchange perfusion system. Whole-cell configuration was first obtained in voltage-clamp mode before proceeding to current-clamp recording mode. Cells with a stable resting membrane potential (RMP) more negative than -40 mV for a minimum of 5 min were used to assess neuronal excitability. Gap-free recordings were made with a 5 min baseline followed by 5 min treatment of either recombinant mouse GM-CSF (300 ng ml^-1^) or vehicle (ECS*_patch_*). Analyzed cells had an average capacitance of ∼15pF, an access resistance < 8 MOhms and an RMP at break-in of ∼53 mV. Cells with an access resistance >10 MOhms and RMP at break-in of >-40mV were excluded. Data were analysed using Clampfit 11.1 (Molecular Devices) with average change in RMP determined by the following formula; *Mean RMP 2 minutes post-drug application – Mean RMP 2 minutes prior to drug application*. Neurons properties are depicted in Supplementary Table 1.

#### DRG neuron multi-electrode array (MEA) electrophysiology

Pooled cryopreserved rat neonatal DRG were purchased from Lonza (Catalog Number: R- DRG-505) and seeded at ∼28,000 cells per well onto Axion Cytoview MEA 12 well plates (Catalog Number: M768-tMEA-12W) as per the manufacturer’s instructions and were maintained by twice weekly feeding (i.e. half media change). Each plate has 768 electrodes shared across 12 wells (64 electrodes per well) and neuronal activity was sampled at 12.5kHz simultaneously from each electrode in each well using the Axion Maestro Pro Multi- Electrode Array instrument. After 14 days in culture, to generate conditioned media each well volume was standardized to 2 ml at least 4 hr prior to experimenting. Recombinant rat GM- CSF or vehicle was then made up to appropriate concentration using 1ml of this conditioned media that was removed from each well. These pre-warmed test substances were then applied and firing behavior monitored at 15 min, 60 min, 4 hr & 24 hr post-treatment. Data was analyzed using the Axion Biosystems AxiS Navigator software and electrodes with ≥ 5 μV baseline noise were excluded from analyses. Spikes were identified with thresholds set at 6x standard deviation of background noise and mean firing rate derived from all recording electrodes per well.

## General experimental design

All *in vivo* experiments were performed in both male and female age-matched mice and littermates when possible. Treatment groups of mice were randomized and evenly distributed across both male and female littermates and/or cagemates. Behavior analyses were performed in a blinded manner. Animal numbers were estimated on the basis of pilot studies or published work.

## Statistical analysis

Unless otherwise stated, data were reported as the means ± s.e.m. The normal distribution of data was analyzed by D’Agostino and Pearson test. Statistical analysis was performed with GraphPad Prism (Version 8.0.1, GraphPad Software, San Diego, CA, USA). Unpaired Student’s t-test was used for 2 group comparisons. For data with multiple groups, one-way ANOVA or 2-way ANOVA was performed followed by *post hoc* corrections which are indicated in each figure legend.

## Code availability

Code for the analysis of scRNA-seq data will be uploaded into a GitHub repository upon publication.

## Data Availability

scRNA-seq data generated from this paper will be made publicly available in a repository upon publication. Additional datasets generated during and/or analyzed during the current study are available from the lead contact on reasonable request. Source data are provided with this paper.

## Acknowledgements

We thank Ieda R. Santos for expert technical assistance in all animal experiments; José Augusto Maulim and Maria Dolores S. Ferreira for scanning electron microscopy experiments; Roberta Ribeiro Costa Rosales and Elizabete Rosa Milani for their support in confocal microcopy acquisition and analyses; Denise Brufato Ferraz for support in flow cytometry data acquisition; Sabrina Baroni for support in scRNAseq data acquisition; Marcella Daruge Grando and Ana Katia dos Santos for managing the laboratory, prepare solution. We also thank the contributions of Jaimini Kumar at GSK for histology support and genOway who developed novel transgenic mouse strains used in the study on behalf of GSK. This work was supported by funding from the São Paulo Research Foundation (FAPESP) under grants agreements n° 2013/08216-2 (Center for Research in Inflammatory Disease), and GSK-UK under grant agreement FMRP-USP/GSK-UK (n. 1011852).

## Author Contributions

G.R.P. performed, participated and analyzed almost all experiments including pain behavioral, flow cytometry, RT-PCR, DRG meninges and frozen tissue for confocal microscopy, human microCT analyses, scRNAseq; prepared all figures included in the manuscript and contributed to the writing and revision of the manuscript. G.V.L.S. performed all bioinformatic analyses and prepared all figures related to these data. W.A.G. established and performed experiments with DRG meninges for confocal microscopy and scanning electron microscopy. D.C.N. S.D. helped in the execution of scRNAseq experiments. G.P.F., M.M.B. helped in microCT analyses in mice. A.U.Q, performed behavioral and confocal microscopy analyses. C.E.A.S. performed immunofluorescence of frozen tissue. J.P.M. Performed the generation of bone marrow-derived macrophages (BMDMs) and ELISA assays. S.E.J. and K.L.M. performed and provided inputs on light-sheet microscopy. V.H. performed light-sheet microscopy, analyses the images and prepared the related figures.

C.G.M and MCC. helped in flow cytometry experiments. T.K.A. performed the experiments related to CCI model, analyzed the data and prepared figures. G.N.R.G., H.I.N. helped in bioinformatic analyses and contributed to data analyses. M.H.N.B. contributed the human tissue specimens and interpreted the human findings. M.M.A. and G.B.M. provided scientific inputs and helped with confocal images acquisition. A.J.D. performed histopathological analyses. J.R.F.H. provided conceptual input, conceived and performed neuronal excitability experiments, analyzed data and interpreted results, prepared figures. E.C.E. provided conceptual input, conceived and coordinated neuronal excitability experiments and development of transgenic mice, interpreted results, secured funding. J.E.S. provided conceptual input, conceived and coordinated neuronal excitability experiments and development of transgenic mice, interpreted results, secured funding. C.D.E. provided conceptual input, conceived and coordinated neuronal excitability experiments and development of transgenic mice, interpreted results, secured funding. F.Q.C., J.C.A.F., acquired funding for this project and contributed to data interpretation and provided conceptual input during the study. T.J.P., I.S. and K.M. re-analyzed spatial transcriptome from human DRG with painful Diabetic Neuropathy, prepared the figure and text related to this data. P.A.G., A.K. and B.B provided scientific and laboratory tools; contributed to writing, review and editing of the manuscript. T.M.C. conceptualized, designed the study, supervised the project and wrote the manuscript. All of the authors contributed to the paper, discussed the results, edited and approved the final manuscript.

## Additional information

Supplementary Information is available for this paper. Correspondence to T.M.C. All request for materials should be addressed to T. M. C.

---

**Extended Data Fig. 1.**
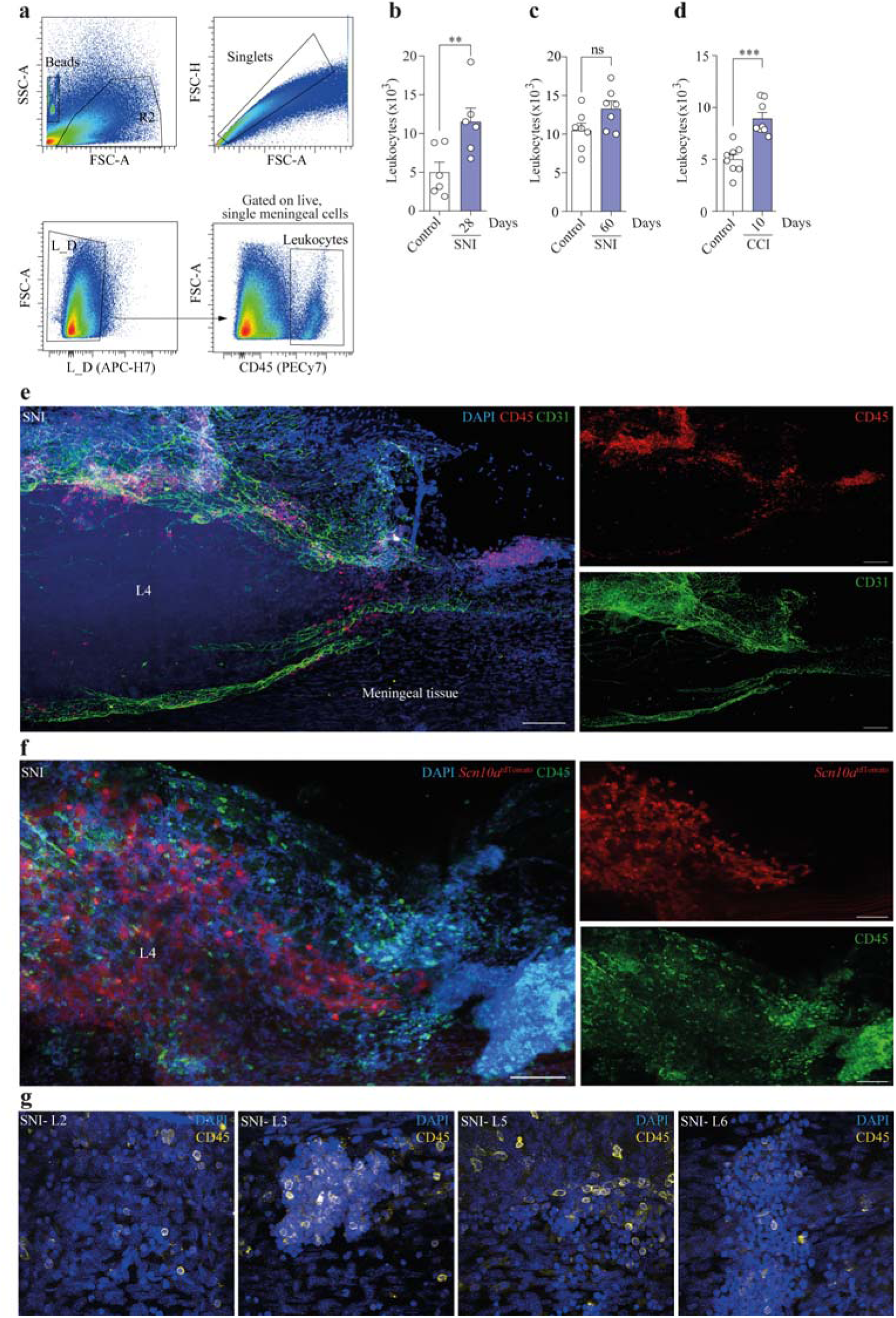
Related to Fig. 1. Leukocytes accumulate in the DRG meninges after peripheral nerve injury. **a.** Flow cytometry gating strategy used to identify leukocyte (CD45^+^ cells) in the DRG meninges from control or SNI mice. **b.** Absolute number of leukocytes (CD45^+^ cells) in the DRG meninges from control or 28 days after SNI. *n* = 6 per group. **c.** Absolute number of leukocytes (CD45^+^ cells) in the DRG meninges from control or 60 days after SNI. *n* = 7 per group. **d.** Absolute number of leukocytes (CD45^+^ cells) in the DRG meninges from control or 10 days after CCI. *n* = 8 per group. **e.** Representative images of meningeal L4 DRG’s whole mounts stained for CD45^+^ cells (red) and CD31^+^ cells (green). DAPI (blue) was used to stain the cell nucleus Samples were harvested from C57Bl/6 mice subjected to SNI (10 days). Scale bar, 100 μm. **f.** Representative images of meningeal L4 DRG’s whole mounts stained for CD45^+^ cells (green) and *Scn10a*^tdTomato^ cells (nociceptive neurons cells body, red). DAPI (blue) was used to stain the cell nucleus. Samples were harvested from *Scn10a*^tdTomato^ reporter mice subjected to SNI (10 days). Scale bar, 100 μm. **g.** Representative images of meningeal L2, L3, L5 and L6 DRG’s whole mounts stained for CD45^+^ cells (yellow). DAPI (blue) was used to stain the cell nucleus Samples were harvested from C57Bl/6 mice subjected to SNI (10 days) or control. Scale bars 50 μm. Statistical analysis: unpaired two-sided *t*-tests (b-d). \*\**P* < 0.01, \*\*\**P* < 0.001. ns = non-significant. n = biologically independent samples from individual mice or mouse tissues. Error bars indicate the mean ± s.e.m.

**Extended Data Fig. 2.**
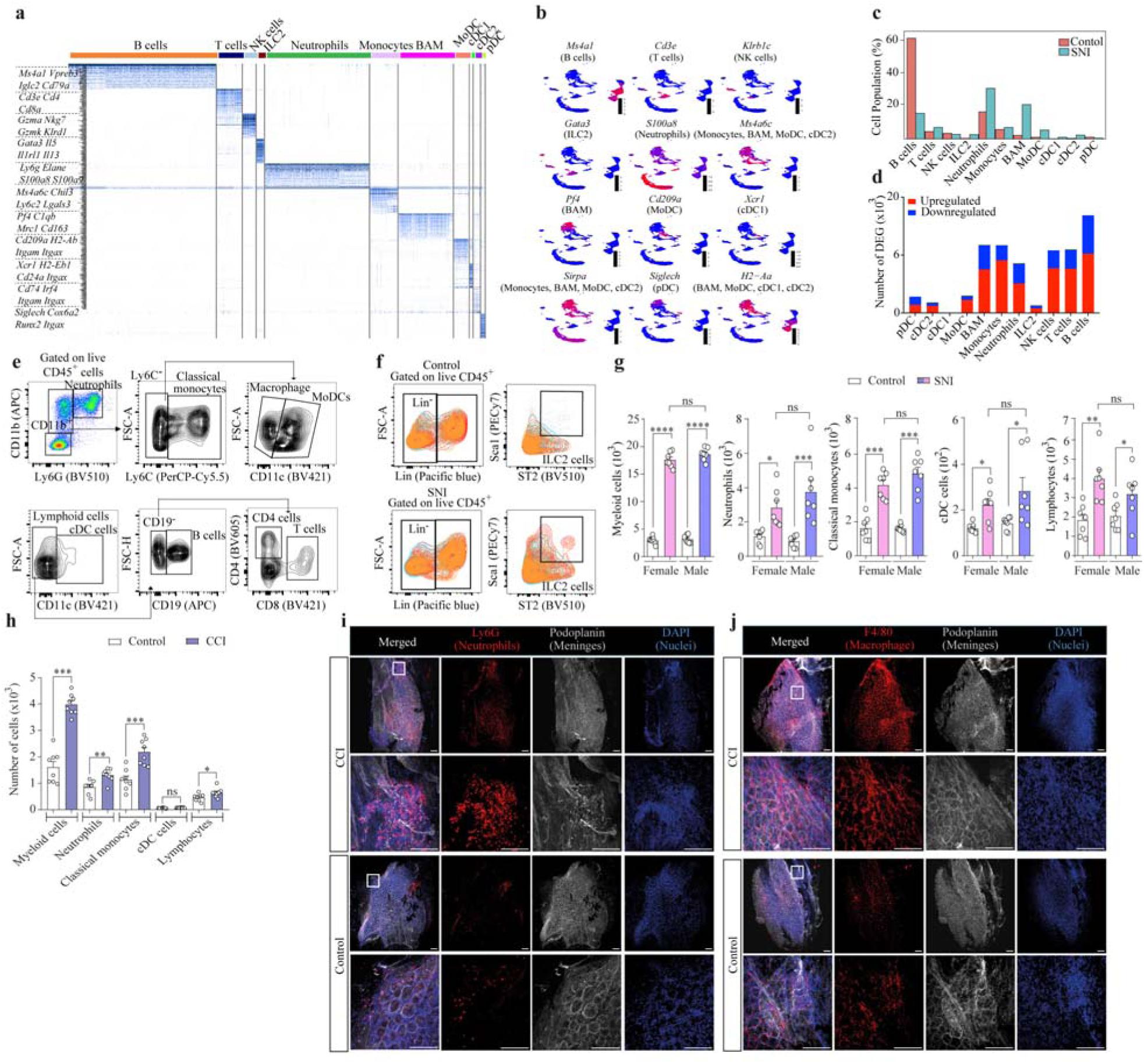
Related to Fig. 2. Single-cell RNA-sequencing analysis of leukocytes from DRG meninges. **a.** Heatmap showing cluster-specific gene expression with unique marker genes used to annotate each cluster. Selected marker genes for major immune cell types are indicated on the left. **b.** Feature plots of representative marker genes for each cluster highlighting their expression patterns across the dataset. **c.** Proportional representation of each cell cluster in control (red) and SNI (green) conditions. **d.** Quantification of differentially expressed genes (DEGs) showing the number of genes upregulated (red) and downregulated (blue) in SNI across each cluster. **e.** Flow cytometry gating strategy to identify major leukocyte populations in the DRG meninges from control or SNI mice. **f.** Flow cytometry gating strategy to identify ILC2s in the DRG meninges from control mice or 10 days after SNI. **g.** Absolute number of different cell populations in the DRG meninges from male *versus f*emale control mice or 10 days after SNI. *n* = 7 per group. **h.** Absolute number of different cell populations in the DRG meninges from control mice or 10 days after CCI. *n* = 8 per group. **i.** Representative images of whole mount preparation containing L4 DRG and attached meninges stained for Ly6G^+^ neutrophils (red), podoplanin^+^ (white, pseudo colored), and DAPI (blue). Samples were harvested from C57Bl/6 mice subjected to CCI (10 days) or control. Scale bars, 100 μm. **j.** Representative images of whole mount preparation containing L4 DRG and attached meninges stained for F4/80^+^ macrophages (red), podoplanin^+^ (white, pseudo colored), and DAPI (blue). Samples were harvested from C57Bl/6 mice subjected to CCI (10 days) or control. Scale bars, 100 μm. Statistical analysis: one-way ANOVA with Tukey post-test (g), unpaired two-sided *t*-tests (h). \**P* < 0.05, \*\**P* < 0.01, \*\*\**P* < 0.001, \*\*\*\**P* < 0.0001. ns = non-significant. *n* = biologically independent samples from individual mice or mouse tissues. Error bars indicate the mean ± s.e.m.

**Extended Data Fig. 3.**
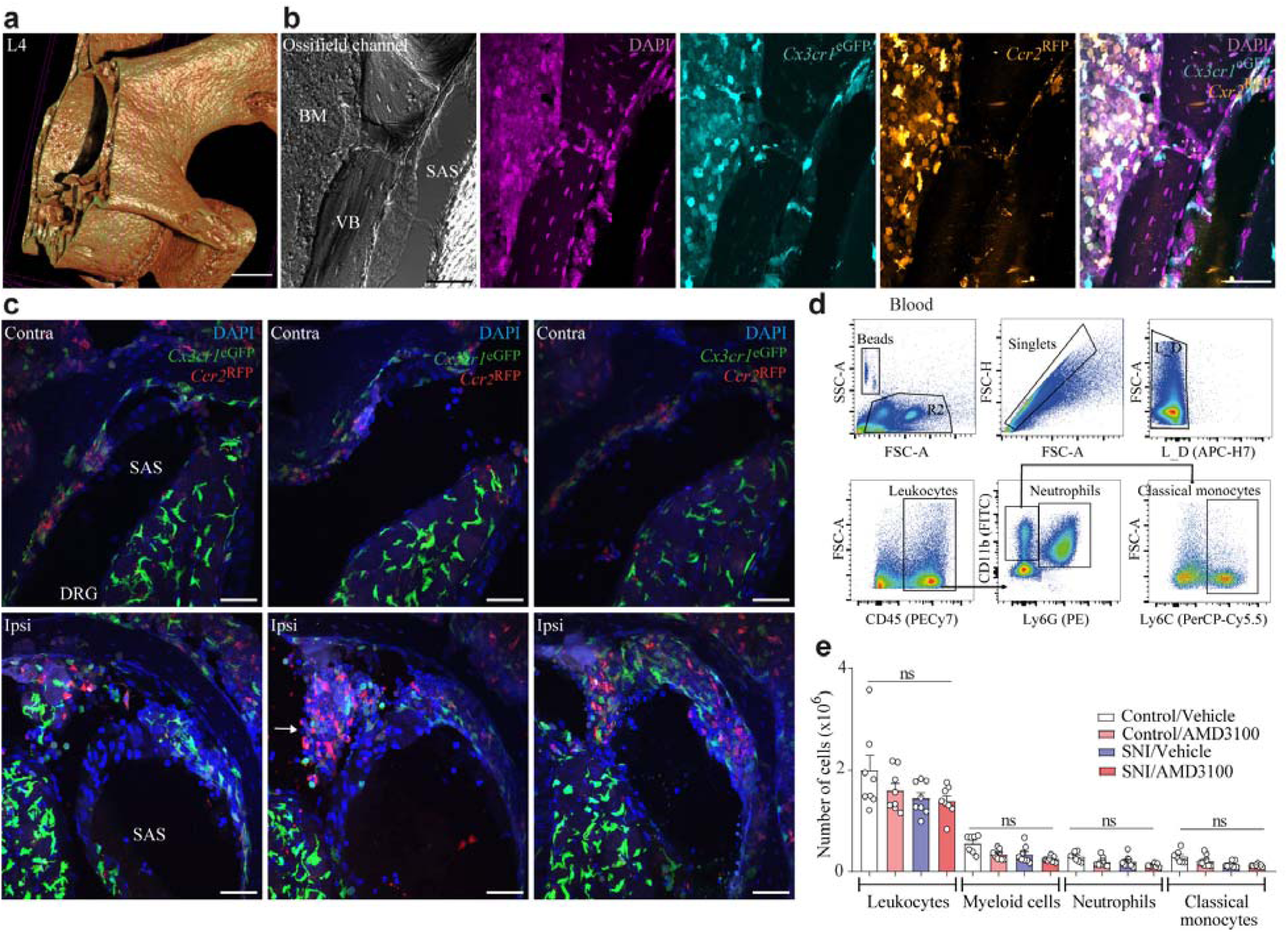
Related to Fig. 3. Myeloid cells migrating from vertebral BM towards DRG’s meninges thought ossified channels. **a.** Digital section showing antero- lateral view of the lumbar vertebra (L4) imaged by micro-CT. Ossified channels openings (dashed circle) on the inner surface of the vertebral bone. Scale bar, 500 μm. **b.** Representative confocal image from maximum intensity projection of lumbar vertebra from *Ccr2*^RFP^ *Cx3cr1*^eGFP^ dual reporter mice. *Ccr2*^RFP^ and *Cx3cr1*^eGFP^ positive leukocytes in the vertebral BM, inside the vertebra bone (VB) ossified channel near the L4 DRG. The confocal image focused on the vertebral arch region showing an ossified channel traversing the VB to connect the vertebral BM to the subarachnoid space (SAS) near the L4 DRG. Sample collection was performed 10 days after SNI surgery in *Ccr2*^RFP^*Cx3cr1*^eGFP^ double transgenic mice. Scale bar, 50 μm. **c.** Confocal image focused on the contralateral (CL) and ipsilateral (Ipsi) region near the dorsal part of L4 DRG. Arrow indicates accumulated cells. Sample collection was performed 10 days after SNI surgery. Scale bar, 50 μm. **d.** Flow cytometry gating strategy to identify total leukocytes (CD45^+^ cells) and myeloid cells (neutrophils and classical monocytes) in the blood of control or SNI mice treated with vehicle or AMD3100. **e.** Absolute number of leukocytes and myeloid cells (neutrophils and monocytes) in the blood of control or SNI mice treated with vehicle or AMD3100 (5 μg/5 μL, i.t.). *n* = 8 per group. Statistical analysis: one-way ANOVA with Tukey post-test (e). ns = non-significant. *n* = biologically independent samples from individual mice or mouse tissues. Error bars indicate the mean ± s.e.m.

**Extended Data Fig. 4.**
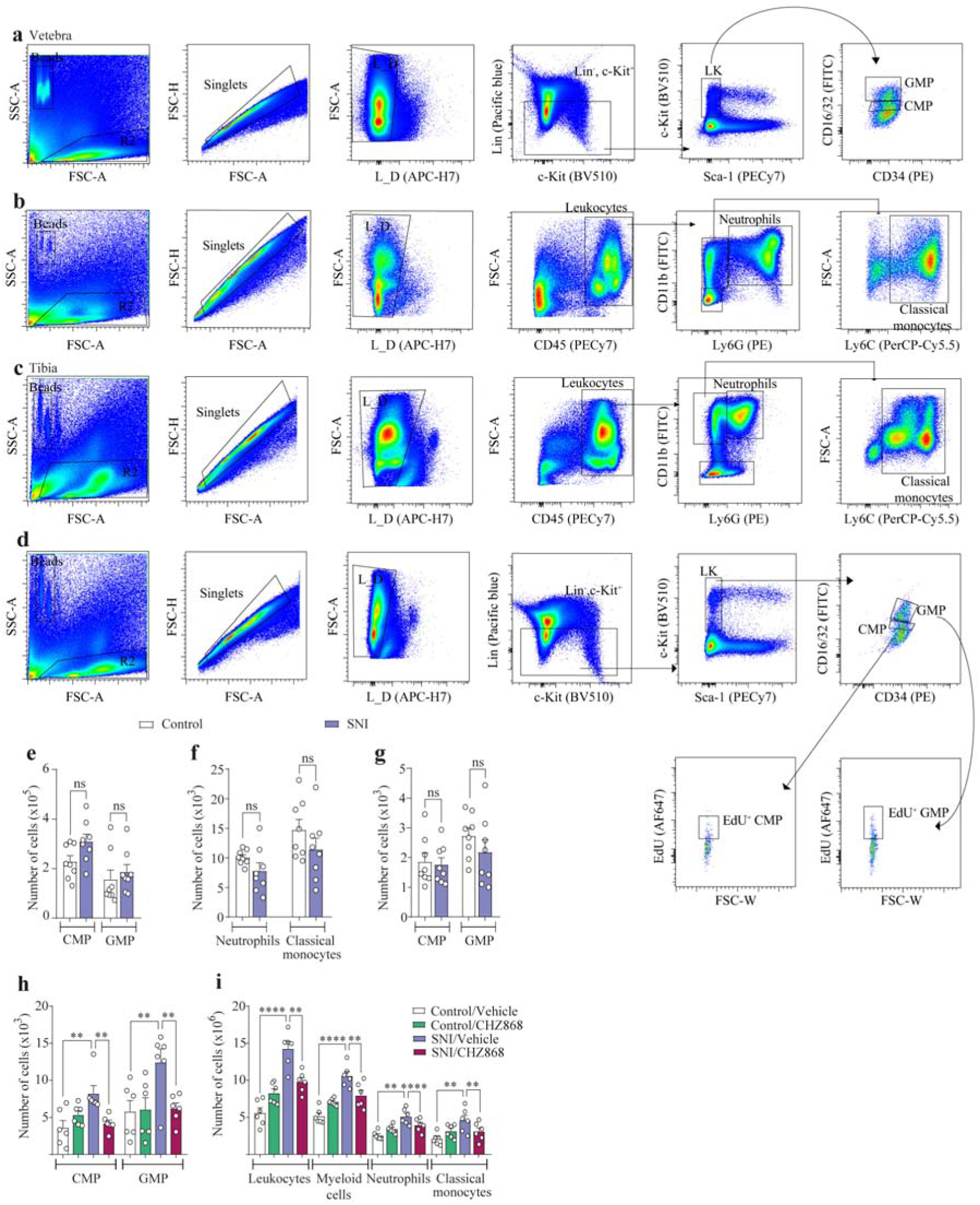
Related to Fig. 4. Emergency myelopoiesis in the vertebral or tibial BM after peripheral nerve injury. **a.** Flow cytometry gating strategy to identify CMP and GMP populations in the vertebral BM from control or SNI mice. **b.** Flow cytometry gating strategy to identify myeloid leukocyte (neutrophils and classical monocytes) populations in the vertebral BM from control or SNI mice. **c.** Flow cytometry gating strategy to identify myeloid leukocyte (neutrophils and classical monocytes) populations in the tibia BM from control or SNI mice. **d.** Flow cytometry gating strategy to identify CMP and GMP progenitor populations in the tibia BM from control or SNI mice. **e.** Absolute number of neutrophils and classical monocytes in the tibia BM from control mice or after 10 days SNI. *n* = 8 per group. **f.** Absolute number of CMP and GMP progenitor populations in tibia BM from control mice or 10 days after SNI. *n* = 8 mice per group. **g.** Absolute number of proliferated EdU^+^CMP and EdU^+^GMP in tibia BM from control mice or 10 days after SNI. *n* = 8 mice per group. **h.** Absolute number of CMPs and granulocyte-monocyte progenitor (GMPs) in the vertebral BM from control mice or 10 days after SNI that receive vehicle or CHZ868 treatments. *n* = 6 per group. **i.** Absolute number of different cell populations in the vertebral BM from control and SNI samples that receive vehicle or CHZ868 treatments. n = 6 per group. Statistical analysis: one-way ANOVA with Tukey post-test (h,i). unpaired two-sided *t*- tests (e-g) \*\**P* < 0.01, \*\*\*\**P* < 0.0001. ns= non-significant. *n* = biologically independent samples from individual mice. Error bars indicate the mean ± s.e.m.

**Extended Data Fig. 5.**
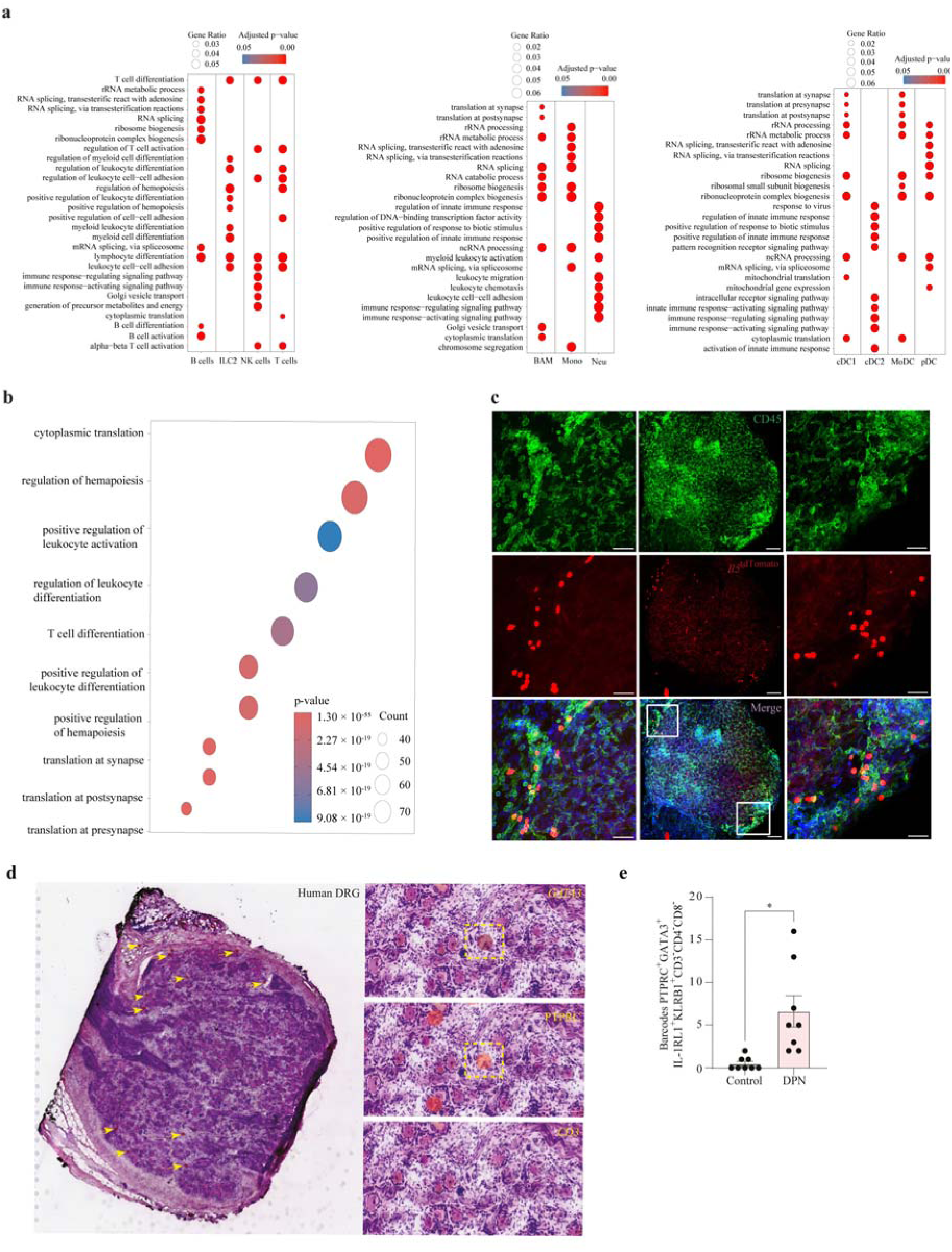
Related to Fig. 5. ILC2s in mouse and human DRG meninges. **a.** Gene Ontology (GO) enrichment analysis of differentially expressed genes in each immune cell cluster using the clusterProfiler package. Dot size represents gene ratio, and color indicates the adjusted p-value for each GO term. **b.** Focused GO enrichment analysis in ILC2 cells highlighting terms related to “regulation of hematopoiesis.” **c.** Representative images of DRG (with meninges) whole-mounts stained for CD45^+^ cells (green), *Il5*^tdTomato^ cells (red, ILC2s) and DAPI (blue). Samples were harvested from *Il5*^tdTomato^ mice subjected to SNI (10 days). Scale bars (from left to right), 100 μm and 50 μm. **d.** Visium spatial transcriptomic data from human DRGs with painful diabetic neuropathy (PDN). Barcodes overlapping with ILC2 markers are indicated by yellow arrows. A representative image shows positive spots for GATA3 and PTPRC and negative for the canonical T-cell marker CD3. **e.** Quantification of barcodes overlapping with ILC2 markers in control versus painful diabetic neuropathy DRGs. *n* = 4 per condition with 2 DRG sections each, and section with 100 μm apart. Statistical analysis: unpaired two-sided *t*-tests (i, j). \**P* < 0.05, Error bars indicate the mean ± s.e.m.

**Extended Data Fig. 6.**
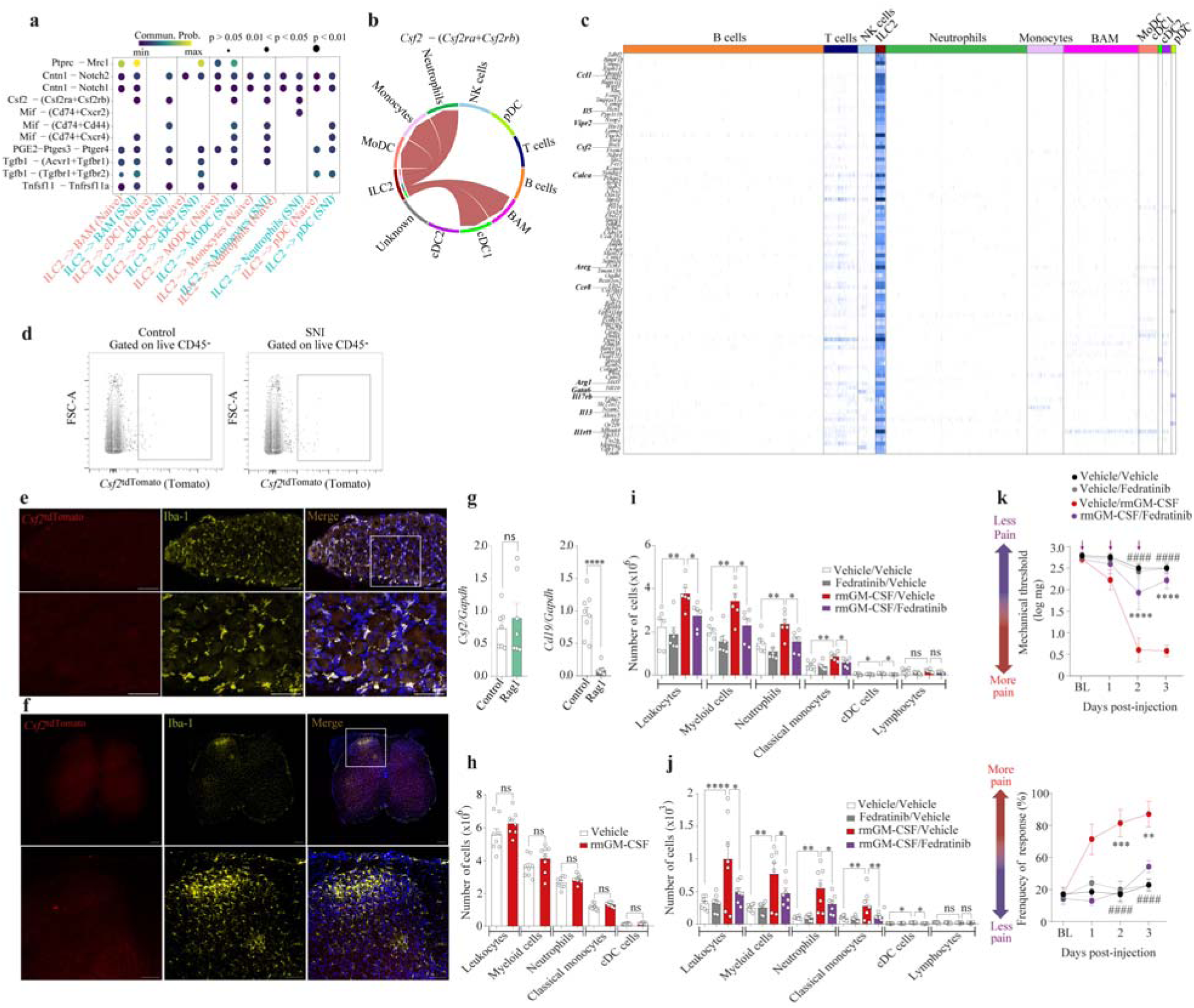
Related to Fig. 5. *Csf2* expressing cells in the DRG meninges and parenchyma and spinal cord. **a.** CellChat analysis of ligand-receptor interactions between ILC2 and other immune populations. This dot plot displays the predicted intercellular communication probabilities between ILC2 and various myeloid cell types under control (pink) and SNI (green) conditions, inferred using CellChat. **b.** Chord diagram of Csf2- mediated outgoing signaling from ILC2. This chord diagram illustrates the predicted outgoing interactions of Csf2 from ILC2 to various immune cell types. Red arcs represent the direction and strength of communication, highlighting robust signaling from ILC2 toward monocytes, MoDC, and neutrophils, suggesting that ILC2s are major source of Csf2 in this context. **c.** Heatmap showing selected key marker genes for ILC2s, including *Ccl1*, *Il5*, *Vipr2*, *Csf2*, *Calca*, *Areg*, *Ccr8*, *Arg1*, *Gata6*, *Il17rb*, *Il13 and Il1rl1* are highlighted on the left. **d.** Flow cytometry gating showing *Csf2*^tdTomato^ cells in CD45^-^ cell populations in samples from the DRG meninges of *Csf2*^tdTomato^ mice. **e.** Representative images showing the expression of Iba1 and *Csf2*^tdTomato^ in the L4 DRG parenchyma. Samples were collected 10 days after induction of SNI in *Csf2*^tdTomato^ mice. Scale bar, (top to bottom) 100 μm and 50 μm. **f.** Representative images showing the expression of Iba1 and *Csf2*^tdTomato^ cells in the lumbar spinal cord. Samples were collected 10 days after induction of SNI in *Csf2*^tdTomato^ mice. Scale bar, (top to bottom) 200 μm and 100 μm. **g.** qPCR analysis of relative expression of *Csf2* and *Cd19* in the DRG meninges from C57BL/6J WT or *Rag1*^−/−^mice. n = 7-8 mice per group. **h.** Absolute number of different cell populations in the tibial BM from mice that received vehicle or rmGM-CSF. *n* = 8 per group. **i.** Absolute number of different cell populations in the vertebral BM from mice treated with vehicle or Fedratinib (60 mg kg^−1^, *p.o*) 1 h before i.t. injection of vehicle or rmGM-CSF (100 ng/site/injection). *n* = 6 per group. **j.** Absolute numbers of different cell populations in the DRG meninges from mice treated with vehicle or Fedratinib (60 mg kg^−1^, *p.o*) 1 h before i.t. injection of vehicle or rmGM-CSF (100 ng/site/injection). *n* = 6 per group. **k.** Analyses of mechanical pain hypersensitivity in *von Frey* test in mice treated with vehicle or Fedratinib (60 mg kg^−1^, *p.o*) 1 h before i.t. injection of vehicle or rmGM-CSF (100 ng/site/injection). *n* = 6 per group. Statistical analysis: two-way ANOVA with Bonferroni post-test (k); one-way ANOVA with Tukey post-test (i,j); unpaired two-sided *t*-tests (g,h). \**P* < 0.05, \*\**P* < 0.01, \*\*\**P* < 0.001, \*\*\*\**P* < 0.0001, ^####^*P* < 0.0001. ns = non- significant. *n* = biologically independent samples from individual mice or mouse tissues. Error bars indicate the mean ± s.e.m.

**Extended Data Fig. 7.**
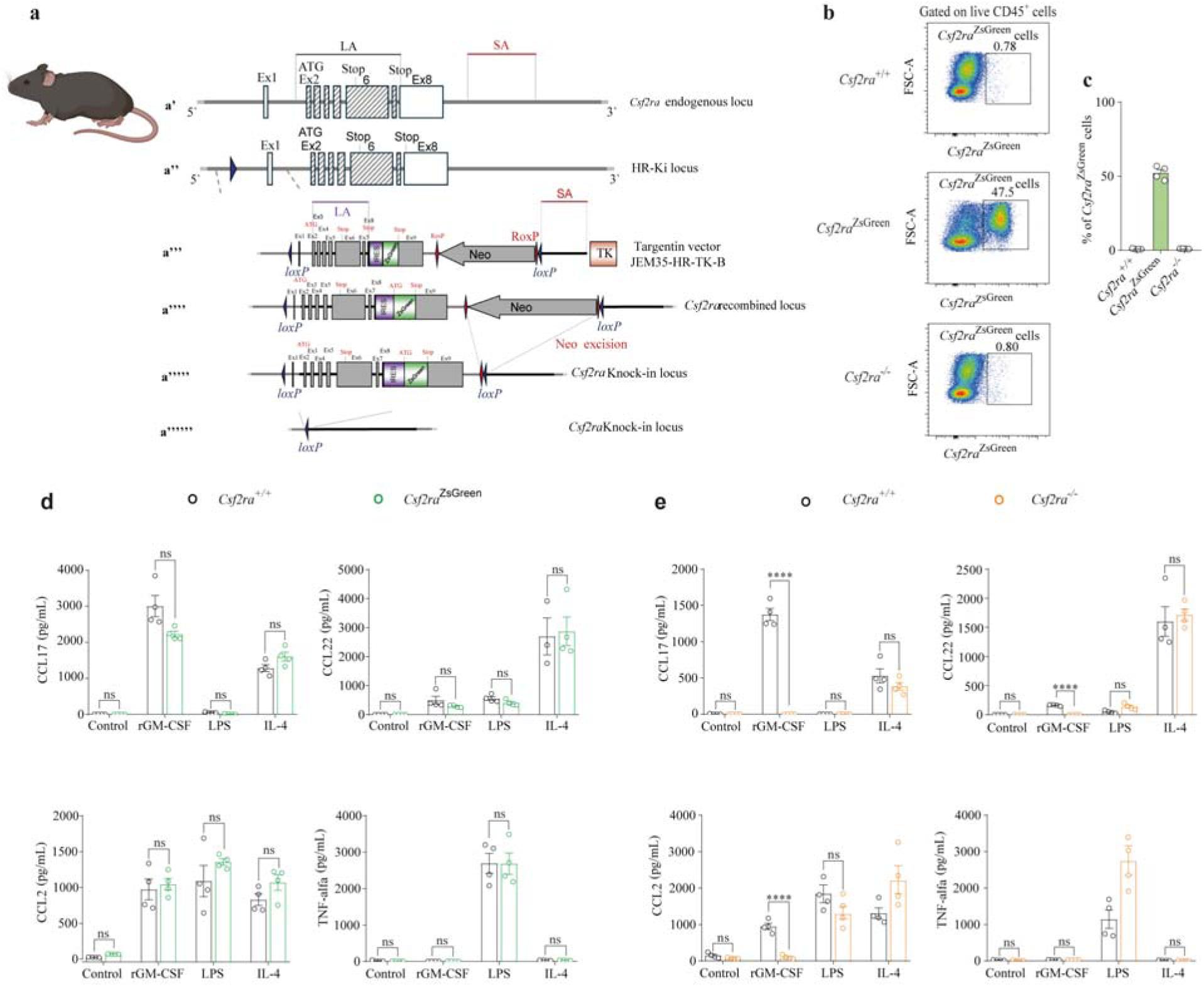
Related to Fig. 6. Generation and functional validation of *Csf2ra*^ZsGreen^ reporter and *Csf2ra^-/-^* mice. **a. a’.** *Csf2ra* endogenous locus. LA, the long homology arm including exons 2-11 and a portion of exon 12 of the NM_009970 transcript. SA, the short homology arm including a 3’ *Csf2ra* region. **a’’.** *Csf2ra* 5’ *loxP* knock-in locus. A *lox*P site (a blue triangle) was inserted in the 5’ region of *Csf2ra* using the CRISPR/Cas9 technology and the homologous recombination (HR) repair pathway in C57BL/6J embryonic stem (ES) cells. The ES cells were transfected with CRISPR/Cas9 components and the single- stranded oligonucleotide (ssODN) coding for the *lox*P site. ES cell clones that had the *lox*P site inserted into both alleles (i.e., homozygous 5’ *loxP* knock-in clones) were characterized by sequencing a region spanning 1kb upstream and 1 kb downstream of the insertion site. The insertion of the ssODN destroyed the Cas9 cutting site, thus avoiding recurrent cleavage of the mutated locus. Three predicted off-target sites (two in intergenic regions, Chr12:118,766,888-118,766,903 and ChrX:7,349,406-7,349,421, and one in the intronic region of *Cdh13*, Chr8:119,277,322-119,277,337) were sequenced in the targeted ES cell clones, and no off-target mutations were detected. **a’’’.** Targeting vector. It consists of the Diphtheria Toxin A chain (DTA)-negative selection marker, LA, IRES-ZsGreen sequence inserted downstream of the STOP codon in exon 12, the remaining portion of the exon 12, the neomycin positive-selection cassette (Neo, a neomycin resistance gene under the control of the pGK promoter) flanked with *roxP* recombination sites (red diamonds), a 3’ *loxP* site (a blue triangle), SA, and the thymidine kinase (TK) negative selection marker. The vector was verified by sequencing. **a’’’’.** Recombined *Csf2ra* locus. The targeting vector was linearized and electroporated into the previously generated recombined ES cell clones in which the 5’ loxP site was inserted in *Csf2ra* locus (shown in panel **b**) according to a proprietary protocol optimized for ES cells transfection. Transfected ES cells were subjected to negative and positive selection. The correct recombination events in neomycin-resistant ES cell clones were verified by PCR and DNA sequence analyses of the entire targeted vector plus at least 1 kb downstream and 1 kb upstream of the homology arms. The number of copies of the neomycin cassette in the genomic DNA of the recombined ES cell clones was assessed using quantitative PCR and has confirmed the presence of a single copy, indicating that no random, non-homologous integration of the targeted vector occurred in the selected clones. To generate chimeras, blastocysts derived from C57BL/6J-*Tyr* albino mice were injected with the characterized ES cell clones and implanted in pseudo-pregnant females that were allowed to develop to term. Chimerism rate of the progeny was assessed by coat color. **a’’’’’.** Csf2ra- ZsGreen-flox locus. To generate the Neo-excised *Csf2ra* allele expressing zsGreen and flanked by the two *loxP* sites, the chimeric founders were bred with C57BL/6J Dre-deleter mice that ubiquitously express the Dre recombinase. Offspring was screened by PCR, and PCR-positive animals were verified by sequencing to confirm removal of the neomycin resistance cassette. **a’’’’’’.** *Csf2ra^-/-^* locus. To generate the knock-out *Csf2ra* allele (*Csf2ra^-/-^*) the chimeric founders were bred with C57BL/6J Cre-deleter mice that ubiquitously express the Cre recombinase. Offspring was screened by PCR, and PCR-positive animals were verified by sequencing to confirm the excision of the floxed region. **b.** Flow cytometry gating and representative plots of *Csf2ra*^Zsgreen^ cells in the vertebrae BM cells. Samples were collected from *Csf2ra*^+/+^, *Csf2ra^ZsGreen^* and *Csf2ra^-/-^* mice. **c.** Frequencies of *Csf2ra*^zsGreen^ cells in the vertebral BM from *Csf2ra*^+/+^, *Csf2ra*^ZsGreen^ and *Csf2ra^-/-^* mice. *n* = 4 per group. **d.** Concentrations of CCL17, CCL22, CCL2 and TNF levels in the supernatant of BMDM originated from *Csf2ra*^+/+^ and *Csf2ra*^ZsGreen^ mice stimulated with recombinant mouse (rm) GM-CSF (20 ng ml^-1^), IL-4 (20 ng ml^-1^) or LPS (100 ng ml^-1^) or vehicle for 24 h. *n* = 4 mice per group. **e.** Concentrations of CCL17, CCL22, CCL2 and TNF in the supernatant of BMDM originated from *Csf2ra*^+/+^ and *Csf2ra^-/-^* mice stimulated with recombinant mouse (rm) GM-CSF (20 ng ml^-1^), IL-4 (20 ng ml^-1^) or LPS (100 ng ml^-1^) or vehicle for 24 h. *n* = 4 mice per group. Statistical analysis: one-way ANOVA with Tukey post-test (d, e). \*\*\*\**P* < 0.0001. ns = non-significant. *n* = biologically independent samples from individual mice or mouse tissues. Error bars indicate the mean ± s.e.m. Illustrations were created with BioRender.com (https://biorender.com).

**Extended Data Fig. 8.**
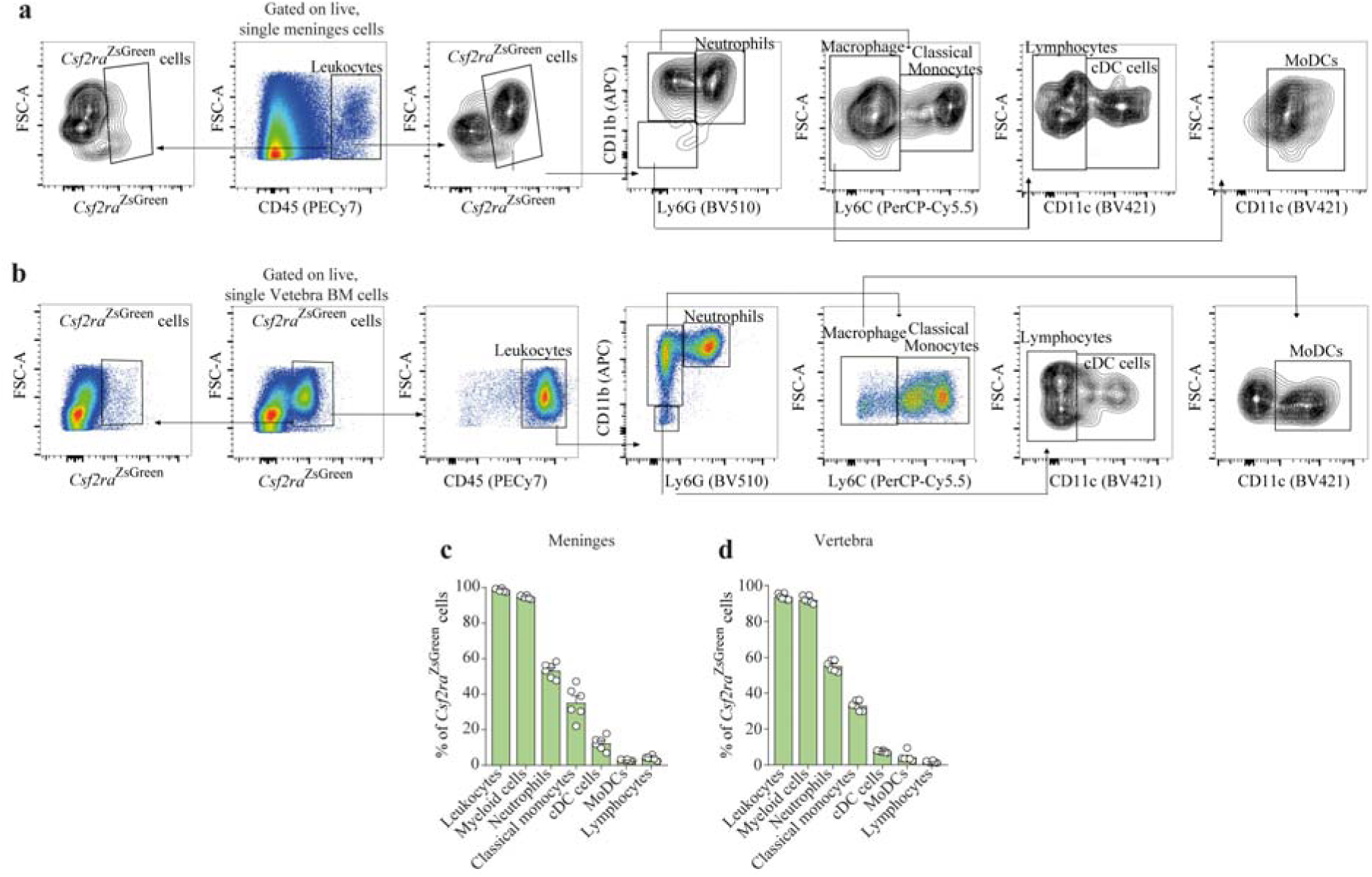
Related to Fig. 6. Flow cytometry analyses of *Csf2ra*^Zsgreen^ cells in the DRG meninges and vertebrae BM. **a.** Flow cytometry gating strategy to identify leukocytes (CD45^+^) and *Csf2ra*^ZsGreen^ cells from DRG meninges. Samples were collected from *Csf2ra*^+/+^ and *Csf2ra*^ZsGreen^ mice. **b.** Flow cytometry gating strategy to identify *Csf2ra*^ZsGreen^ cells from vertebral BM cells. Samples were collected from Csf2ra^+/+^ and *Csf2ra*^ZsGreen^ mice. **c.** Relative frequencies of *Csf2ra*^ZsGreen^ cells within immune populations in the DRG meninges. *n* = 6 mice. **d.** Relative frequencies of *Csf2ra*^ZsGreen^ cells within immune populations in the vertebrae BM. *n* = 6 mice.

**Extended Data Fig. 9.**
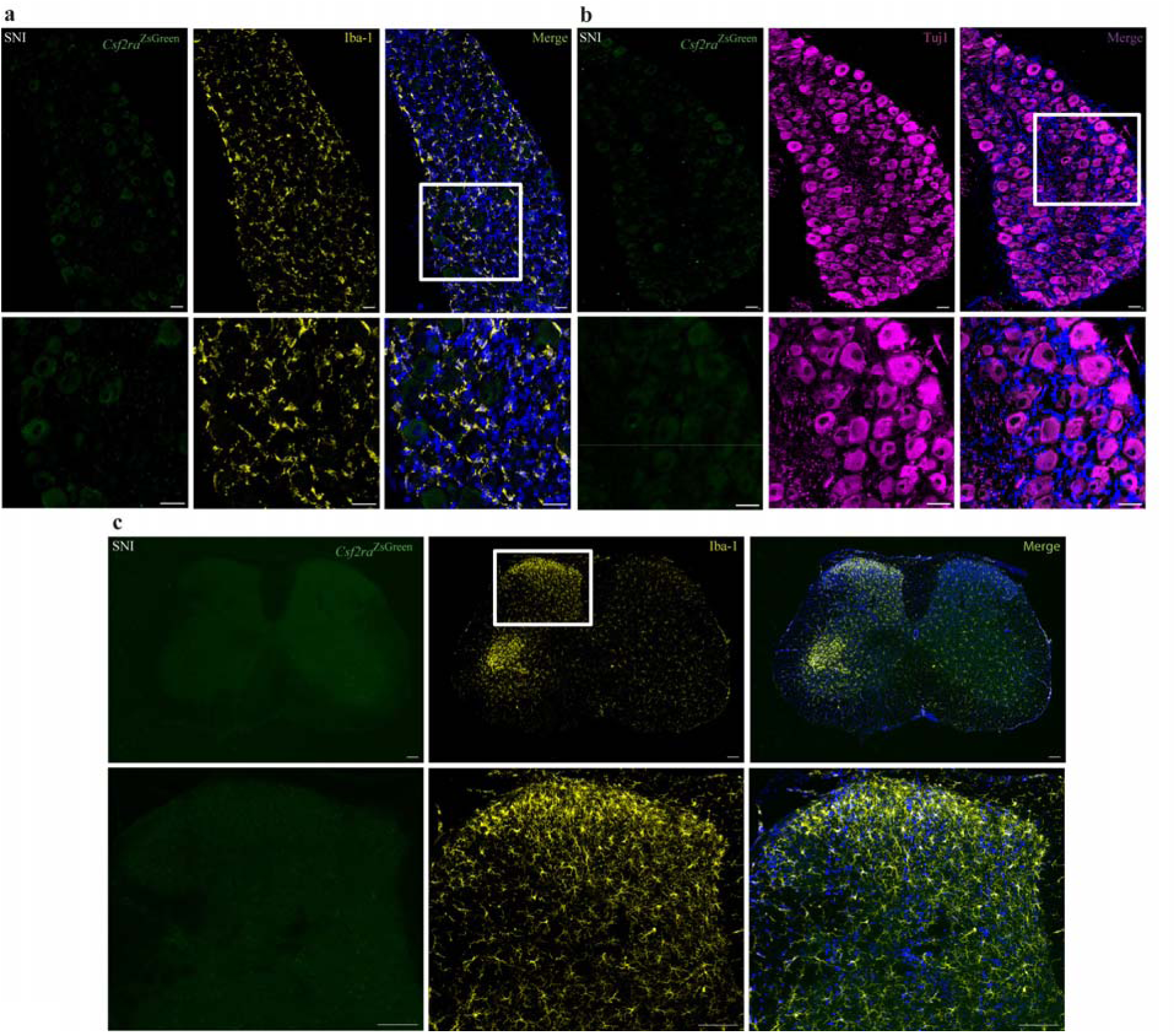
Related to Fig. 6. Lack of *Csf2ra*^Zsgreen^ cells in the DRG and spinal cord parenchyma. **a.** Representative images showing the expression of Iba1 (yellow), ZsGreen (green) in the L4 DRG sections. DAPI stained the cell nuclei. Samples were collected 10 days after induction of SNI in *Csf2ra*^ZsGreen^ mice. Scale bars (from top to below), 100 μm and 50 μm. **b.** Representative images showing the expression of *Tuj1 (*magenta) and ZsGreen (green) in the L4 DRG sections. DAPI stained the cell nuclei. Samples were collected 10 days after induction of SNI in *Csf2ra*^ZsGreen^ mice. Scale bars (from top to below), 100 μm and 50 μm. **c.** Representative images showing the expression of Iba1 (yellow) and ZsGreen (green) in the spinal cord sectins. DAPI stained the cell nuclei. Samples were collected 10 days after induction of SNI in *Csf2ra*^ZsGreen^ mice. Scale bars (from top to below), 200 μm and 100 μm.

**Extended Data Fig. 10.**
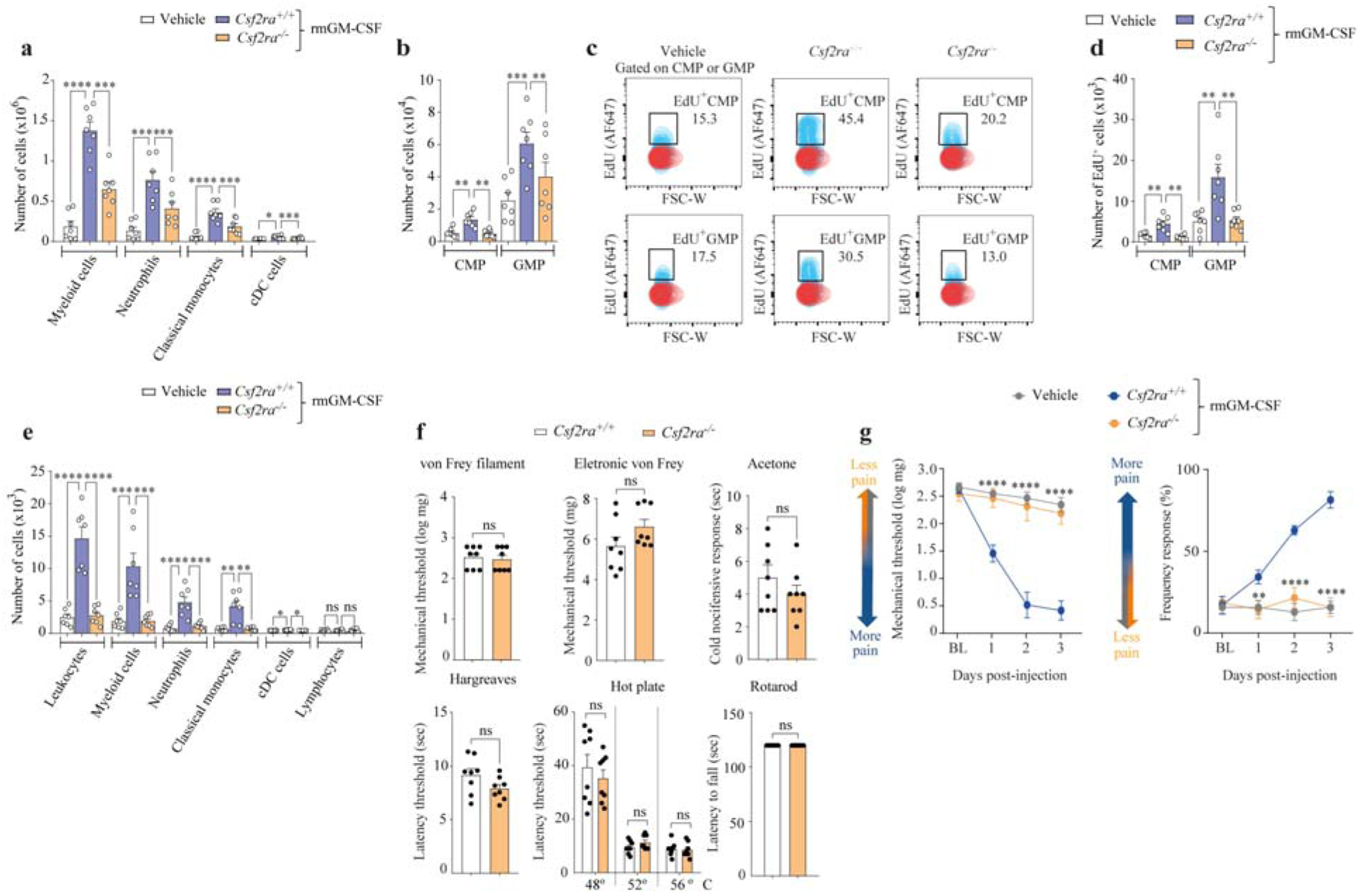
Related to Fig. 6. Meningeal activities of GM-CSF depends on CSF2RA signaling. **a.** Absolute number of different cell populations in vertebral BM from *Csf2ra*^+/+^ and *Csf2ra*^-/-^ mice that received i.t. injection of vehicle or rmGM-CSF (100 ng/site/injection). *n* = 7 per group. **b.** Absolute number of CMP and GMP cells in the vertebral BM from *Csf2ra*^+/+^ and *Csf2ra*^-/-^ mice that received i.t. injection of vehicle or rmGM-CSF (100 ng/site/injection). *n* = 7 per group. **c.** Representative flow cytometry gating plots showing EdU^+^CMP and EdU^+^GMP cells in the vertebral BM. **d.** Absolute number of proliferated (EdU^+^) CMP and EdU^+^GMP cells in the vertebral BM from *Csf2ra*^+/+^ and *Csf2ra*^- /-^ mice that received i.t. injection of vehicle or rmGM-CSF (100 ng/site/injection). *n* = 7 per group. **e.** Absolute number of different cell populations in the DRG meninges from *Csf2ra^+/+^*and *Csf2ra^-/^*^-^ mice that received i.t. injection of vehicle or rmGM-CSF (100 ng/site/injection). *n* = 7 per group. **f.** Physiological nociceptive and motor behaviors in *Csf2ra*^+/+^ and *Csf2ra*^-/-^ mice. *n* = 8 per group. **g.** Mechanical pain hypersensitivity before (baseline, BL) and 6 hours after each i.t. injection of vehicle or rmGM-CSF (100 ng/site/injection) in *Csf2ra*^+/+^ and *Csf2ra*^-/-^ mice. *n* = 7 per group. Statistical analysis: two-way ANOVA with Bonferroni post-test (g); one-way ANOVA with Tukey post-test (a,b,d,e); unpaired two-sided *t*-tests (f). \**P* < 0.05, \*\**P* < 0.01, \*\*\**P* < 0.001, \*\*\*\**P* < 0.0001. ns = non-significant. *n* = biologically independent samples from individual mice or mouse tissues. Error bars indicate the mean ± s.e.m.

**Extended Data Fig. 11.**
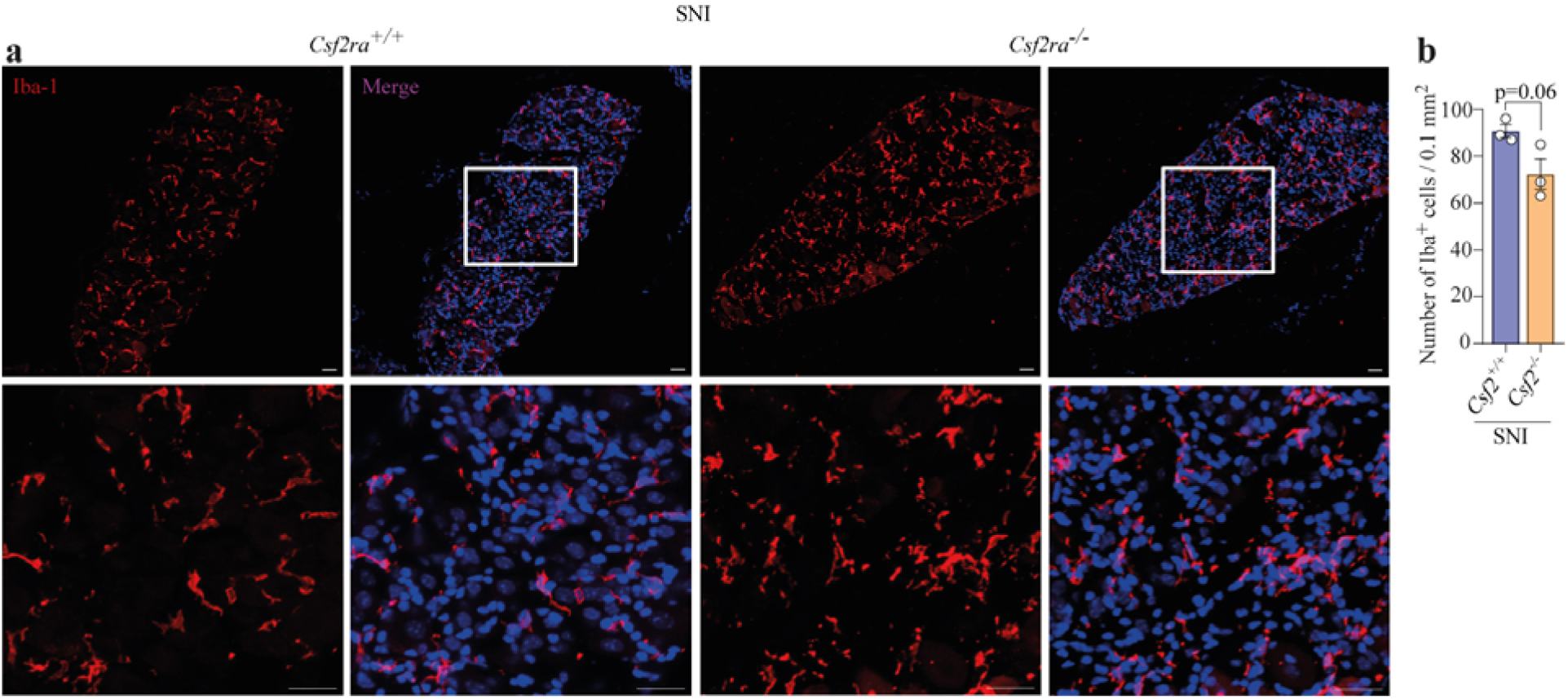
Related to Fig. 6. CSF2RA signaling is not involved in DRG macrophages activation after SNI. **a.** Representative images showing the expression of Iba1 (red) in the L4 DRG sections. DAPI stained the cell nuclei. Samples were collected 10 days after induction of SNI in *Csf2ra^+/+^* or *Csf2ra^-/-^* mice. Scale bars (from top to below), 100 μm and 50 μm. **b.** Quantification of the number of Iba1-positive cells in the L4 DRGs from *Csf2ra^+/+^*or *Csf2ra^-/-^* mice 10 days after SNI. *n* = 3 mice per group. Statistical analysis: unpaired two-sided *t*-tests (b). ns = non-significant. *n* = biologically independent samples from individual mouse tissues. Error bars indicate the mean ± s.e.m.

**Extended Data Fig. 12.**
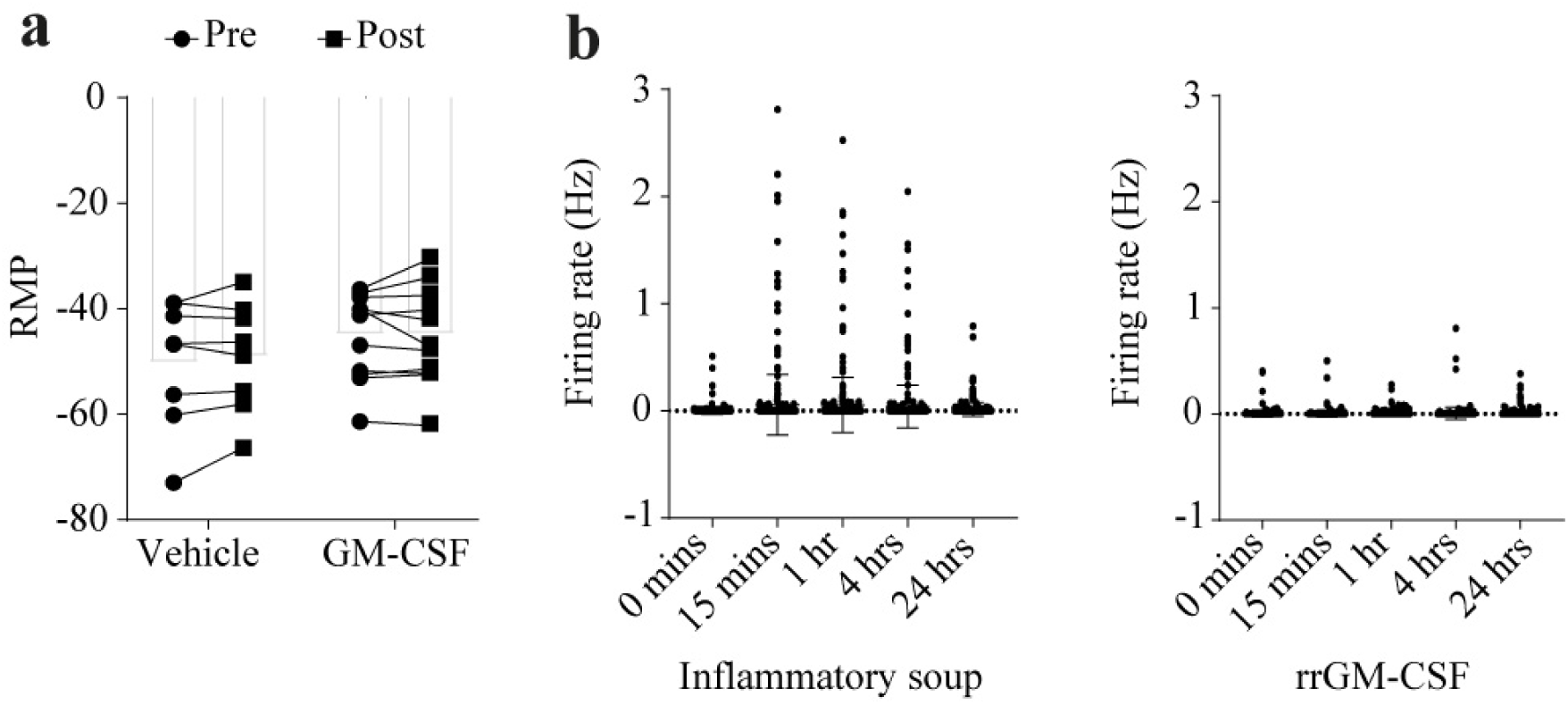
Related to Fig. 6. Effect of GM-CSF on excitability of cultured DRG neurons. **a.** Resting membrane potential (RMP) of cultured mouse DRG neurons before and after the application of recombinant mouse GM-CSF (rmGM-CSF 100 ng ml^-1^; *n* = 8 cells, *N* = 4 independent experiments) or vehicle (*n* = 11 cells, *N* = 5 independent experiments). **b.** Multielectrode (MEA) recordings of neuronal firing rate (Hz) in neonatal rat DRG neurons at different time points over a 24-hour application of recombinant rat GM-CSF (rrGM-CSF 100 ng ml^-1^; *n* = 5 wells, *N* = 2 independent experiments, Number of electrodes recorded: 321) or inflammatory soup (*n* = 6 wells, *N* = 1 independent experiment, Number of electrodes recorded: 384). Statistical analysis: unpaired two-sided t-test (a) vehicle vs. GM-CSF, t(17) = 0.8257, *P* = 0.4204, one-way ANOVA with Dunnett’s post-hoc (b). vehicle: vs. GM-CSF p=0.99, vehicle vs. IS *P* =0.0001. Error bars indicate the mean ± s.e.m.

**Extended Data Fig. 13.**
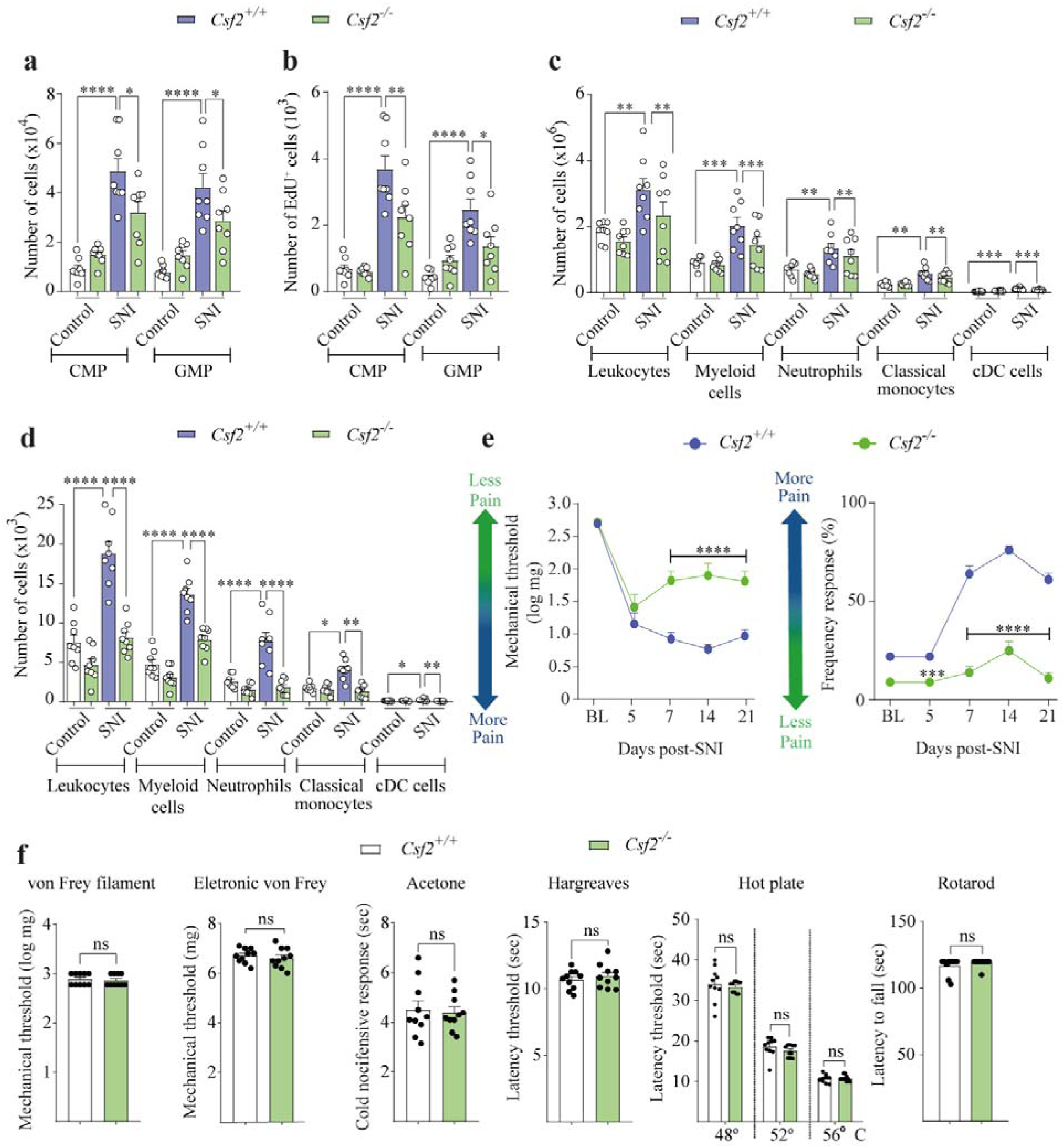
Related to Fig. 6. GM-CSF mediates neuropathic pain and vertebrae bone emergency myelopoiesis after peripheral nerve injury. **a.** Absolute number of common myeloid progenitors (CMPs) and granulocyte-monocyte progenitor (GMPs) in the vertebral BM from *Csf2*^+/+^ or *Csf2*^-/-^ mice control or 10 days after SNI. *n* = 8 per group. **b.** Absolute number of proliferated EdU^+^CMP and EdU^+^GMP in the vertebral BM from *Csf2*^+/+^ or *Csf2*^-/-^ mice control or 10 days after SNI. *n* = 8 per group. **c.** Absolute number of different cell populations in the vertebral BM from *Csf2*^+/+^ or *Csf2*^-/-^ mice control or 10 days after SNI. *n* = 8 per group. **d.** Absolute number of different cell populations in the DRG meninges from *Csf2*^+/+^ or *Csf2*^-/-^ mice control or 10 days after SNI. *n* = 8 per group. **e.** Time course of mechanical pain hypersensitivity in *Csf2*^+/+^ or *Csf2*^-/-^ mice before (baseline, BL) and after SNI. *n* = 8 per group. **f.** Physiological nociceptive and motor behaviors in *Csf2*^+/+^ and *Csf2*^-/-^ mice. *n* = 10 per group. Statistical analysis: two-way ANOVA with Bonferroni post- test (e); one-way ANOVA with Tukey post-test (a-d); unpaired two-sided *t*-tests (f). \**P* < 0.05, \*\**P*< 0.01, \*\*\**P* < 0.001, \*\*\*\**P* < 0.0001. ns = non-significant. *n* = biologically independent samples from individual mice or mouse tissues. Error bars indicate the mean ± s.e.m.

**Extended Data Fig. 14.**
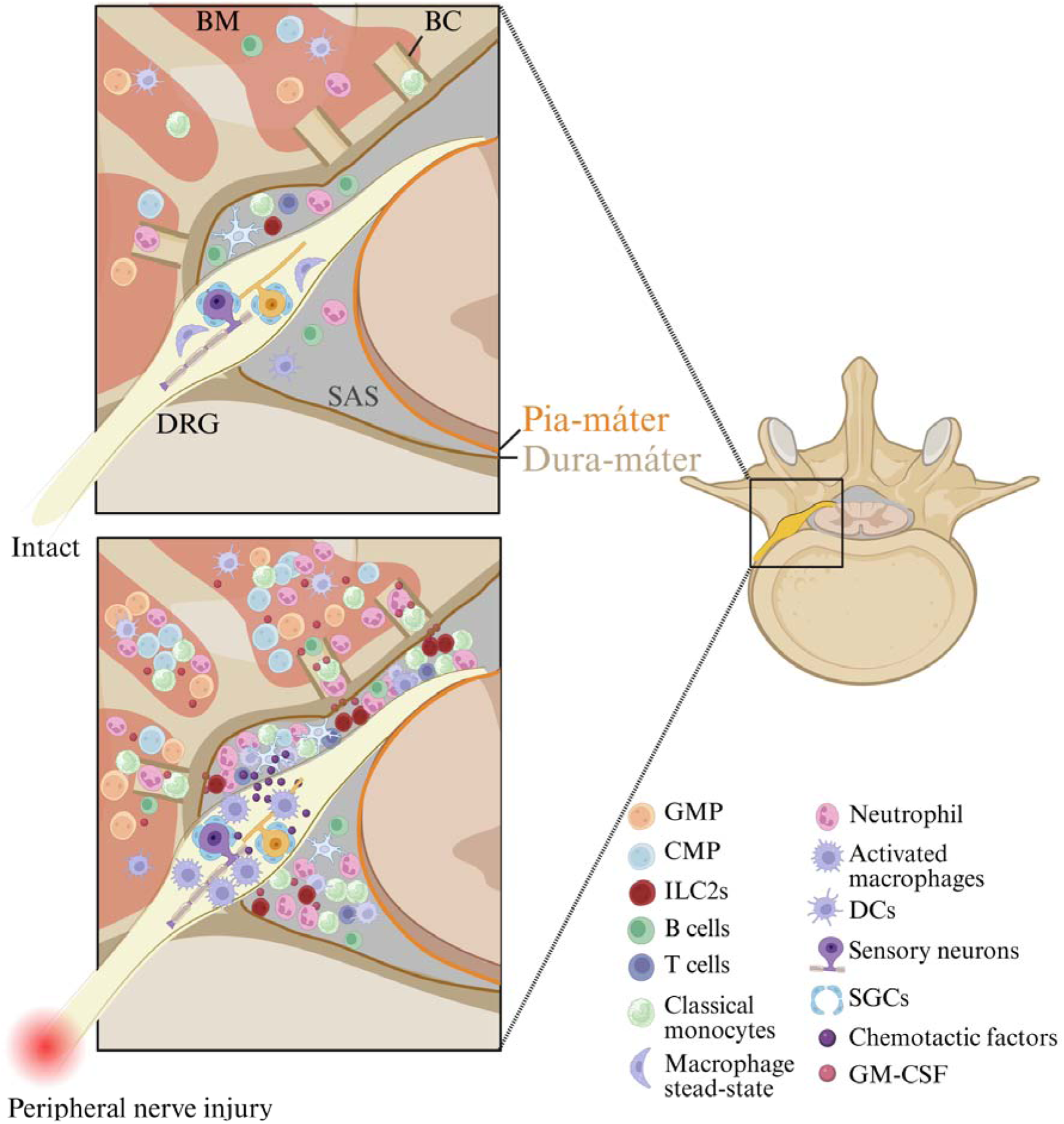
Schematic representation of DRG meninges-vertebral BM axis in naïve condition and during peripheral nerve injury-induced neuropathic pain. After peripheral nerve injury, GM-CSF is released mainly from meningeal ILC2 cells, and can reach the vertebrae BM to induce emergency myelopoiesis. *De novo* produced myeloid cells can directly migrate from vertebral BM toward the DRG meninges through vertebral bone channels. In the last instance, these cells contribute to the development of peripheral nerve injury-induced neuropathic pain. Illustrations were created with BioRender.com (https://biorender.com).

**Extended Data Table 1.** Electrophysiological properties of sensory neurons after GM-CSF treatment.

| Treatment | Capacitance (pF) |  |  | Membrane resist. (MOhm) |  |  | Access resist. (MOhm) |  |  | Holding current (pA) |  |  |
| --- | --- | --- | --- | --- | --- | --- | --- | --- | --- | --- | --- | --- |
|  | Mean | s.e.m. | n | Mean | s.e.m. | n | Mean | s.e.m. | n | Mean | s.e.m. | n |
| Vehicle | 15.87 | 1.761 | 8 | 462.1 | 167.8 | 8 | 7.213 | 1.67 | 8 | -33.81 | 17.48 | 8 |
| GM-CSF<br>100 ng ml<sup>-1</sup> | 15.26 | 0.7078 | 11 | 425.3 | 58.04 | 11 | 7.682 | 0.8948 | 11 | -42.97 | 15.0 | 11 |

Values represent the mean ± s.e.m.; n indicates number of recorded cells.

**Supplementary Video 1: Transverse view of leukocytes in the vertebral column.**

Light-sheet fluorescence microscopy of the cleared vertebral column showing CD45^+^ immune cells (green) distributed within the meninges and surrounding tissues. The sample was optically cleared and immunostained for CD45 to visualize leukocyte localization in three dimensions. BM: bone marrow, DRG: dorsal root ganglion, SC: spinal cord.

**Supplementary Video 2: Medio-lateral view of leukocytes in the vertebral column.**

Light-sheet fluorescence microscopy of the cleared vertebral column showing CD45^+^ immune cells (green) distributed within the meninges and surrounding tissues. The sample was optically cleared and immunostained for CD45 to visualize leukocyte localization in three dimensions. DRG: dorsal root ganglion.

**Supplementary Video 3: Dorso-ventral view of leukocytes in the vertebral column.**

Light-sheet fluorescence microscopy of the cleared vertebral column showing CD45^+^ immune cells (green) distributed within the meninges and surrounding tissues. The sample was optically cleared and immunostained for CD45 to visualize leukocyte localization in three dimensions. DRG: dorsal root ganglion.

